# Ketogenic interventions prevent alterations of the gut microbiome in transgenic Alzheimer’s Disease mice

**DOI:** 10.1101/2025.02.25.640129

**Authors:** Paule E. H. M’Bra, Mariano Avino, Evelyne Ng Kwan Lim, Marian Mayhue, Philippe Balthazar, Anne Aumont, Karine Prévost, Eric Massé, Karl J.L. Fernandes

**Affiliations:** Research Center on Aging, CIUSSS de l’Estrie-CHUS, Sherbrooke, Canada; Department of Medicine, Faculty of Medicine and Health Sciences, Université de Sherbrooke, Sherbrooke, Canada; Department of Neurosciences, Faculty of Medicine, Université de Montréal, Montreal, Canada; Bioinformatic platform, Faculty of Medicine and Health Sciences, Université de Sherbrooke, Sherbrooke, Canada; Department of Biochemistry and Functional Genomics, Université de Sherbrooke, Sherbrooke, Canada

**Author notes:** Correspondance to: Eric Massé, PhD, Karl Fernandes, PhD.

**Keywords:** Alzheimer’s Disease, Microbiome, Medium-chain triglycerides, Ketones

## Abstract

Alterations in the gut microbiome constitute a feature of aging and therefore represent a therapeutic target for aging-related diseases. In this study, we investigated the impact of ketogenic interventions on the microbiome of mice genetically predisposed to Alzheimer’s disease (AD). AD mice exhibited several microbial alterations, notably increased levels of *Bifidobacterium* and decreased levels of *Bacteroidetes.* Ketogenic interventions, either a medium-chain triglyceride-enriched diet (MCT) or carbohydrate-free high-fat diet (CFHF), administered for 1 month restored the levels of more than 50% of the bacteria altered in AD mice, including a strong reduction in *Bifidobacterium* levels. Ketogenic interventions induced a shift in the gut microbiome associated with increased levels of short-chain fatty acid-producing bacteria, such as *Lachnospiraceae* and *Muribaculaceae.* MCT and CFHF also triggered diet-specific microbial changes, which may contribute to the distinct physiological effects of these diets. In conclusion, ketogenic interventions may influence AD pathophysiology by modulating the gut microbiome.

## INTRO

Alzheimer’s Disease (AD) is characterized by pathological changes in the brain (amyloid plaques, neurofibrillary tangles, metabolic defects, and neuroinflammation) that lead to neuronal loss and progressive cognitive decline^1–5^. The exact pathogenesis of AD is not well understood. Cases of sporadic or late-onset AD (representing 95% of all AD cases) are multifactorial and influenced by environmental risk factors that first impair the peripheral system^4,6^. This suggests a key role for body-brain interactions in the etiology of AD. Lifestyle interventions affecting peripheral metabolism, such as ketogenic dietary strategies, yield promising results in reducing mild cognitive deficits^7^ and improving autonomy in patients with AD^8^. The response to such interventions is variable, with some non-responding patients. Understanding the mechanisms mediating the cognitive effects of ketogenic interventions could provide new ideas for optimizing these interventions and counteracting AD. Ketone bodies, such as acetoacetate and beta-hydroxybutyrate (BHB), constitute a cell-permeable energy source for the brain that is preserved during aging and AD. The classic ketogenic intervention is a low-carbohydrate, high-fat diet, which triggers a metabolic switch that boosts ketone production through increased hepatic oxidation of endogenous fatty acids ^9^. Because of the difficulty in adhering to such a diet for a long time, the use of alternatives such as dietary supplementation with ketogenic medium-chain triglycerides (MCT)^10^ made of capric and caprylic acids^11,12^ has been explored. Thus, the most studied mechanisms of action of ketogenic interventions are their effect on improving brain energy metabolism and reducing glucose and body fat metabolic defects, both associated with AD pathophysiology and AD risk^13–15^.

One aspect of ketogenic interventions that remains poorly studied is their indirect and/or non-energetic effects^16,17^, such as their impact on remodeling of the gut microbiome, a feature increasingly considered in AD pathophysiology^17–22^. The gut contains trillions of microorganisms that contribute to intestinal metabolic and immune functions. The microbial abnormalities observed in AD patients appear to be correlated with core features of the disease^19,23,24^ and are thought to promote blood-brain barrier and intestinal barrier permeabilization, increase peripheral and central inflammation, and alter various metabolite levels^25,26^. Microbial metabolites mediate the downstream effects of microbiome alterations. Among them, short-chain fatty acids (SCFA) are the most studied in AD for their potential effects on amyloidogenesis, inflammation, and neuroactive molecules^27–29^. SCFA such as acetate, propionate, and butyrate have one to six carbons in their carbon chain and are produced from the fermentation of dietary fibers in the colon^28,30^. SCFA-producing bacteria, including *Ruminococcus, Bacteroides, Lachnospiraceae (Roseburia, Blautia), Clostridium, Atopobium, and Eubacterium,* differ in the substrate used and the type of SCFA they produce^30,31^. Lactate-producing bacteria, such as *Bifidobacterium* and *Lactobacillus*, can also produce SCFA under certain conditions^30,31^. SCFA levels and SCFA-producing bacteria appear to decrease with aging and AD^31,32^, but the causes of such defects are not well identified, given that the microbiome is highly influenced by diet and genetics. In low-carbohydrate ketogenic diets, ketone bodies absorbed in the gut can serve as intermediates in SCFA metabolic processes bypassing SCFA production from carbohydrates^33^. Nonetheless, the reported effects of ketogenic interventions on SCFA levels and SCFA-producing bacteria are controversial and require further investigations^34–36^.

In this study, we report baseline microbial changes occurring in mouse models of AD and analyzed the changes appearing after ketogenic interventions.

## RESULTS

### Monitoring of metabolic parameters and faecal microbiome features in pre-symptomatic AD mice treated with ketogenic interventions

The gut microbiome is influenced by diet and genetics. Here, we investigated the effects of dementia-causing genetic mutations and ketogenic interventions on the mouse microbiome. The genetic mutations in the 5xFAD model trigger appearance of cerebral amyloid plaques, yet without cognitive deficits at the age we tested (from 2 months old to 5 months old)^37^. The 3xTg-AD model develops amyloid plaques, neurofibrillary tangles, and cognitive deficits later^38^ than the tested period (3 months old to 8 months old). This model also develops peripheral alterations that appeared during the test period. Owing to differences in the manifestation of AD features between the two models, different time points were tested in this study.

3xTg-AD mice, 5xFAD mice, and their wild-type aged-match counterparts (WT) were fed either a control diet (70% carbohydrates, 20% long-chain triglycerides, and 10% proteins, 4.1 kcal/g), an MCT diet (70% carbohydrates, 10% long-chain triglycerides, 10% medium-chain triglycerides, and 10% proteins, 4.1 kcal/g), or a CFHF diet (90% long-chain triglycerides and 10% proteins, 6.7 kcal/g), with equal micronutrient proportion. First, we analyzed the physiological response of mice to ketogenic interventions by measuring body weight and beta-hydroxybutyrate, the main circulating ketone body.

3xTg-AD mice receiving the CFHF diet for 5 months gained twice as much weight as WT mice fed the same diet (**Fig.1a**), showing a sustained increase in blood ketone levels that was nevertheless dampened compared with that in WT mice (**Fig.1b**). Mice fed MCT did not show changes in body weight or blood ketone levels changes (**Fig.1a-b**). 5xFAD mice on the CFHF diet also presented higher body weight gain (**Supp.Fig.1a**) and elevated blood ketone levels with a dampened increase at D85 compared to their WT counterparts (**Supp.Fig.1b**).

**Figure 1.**
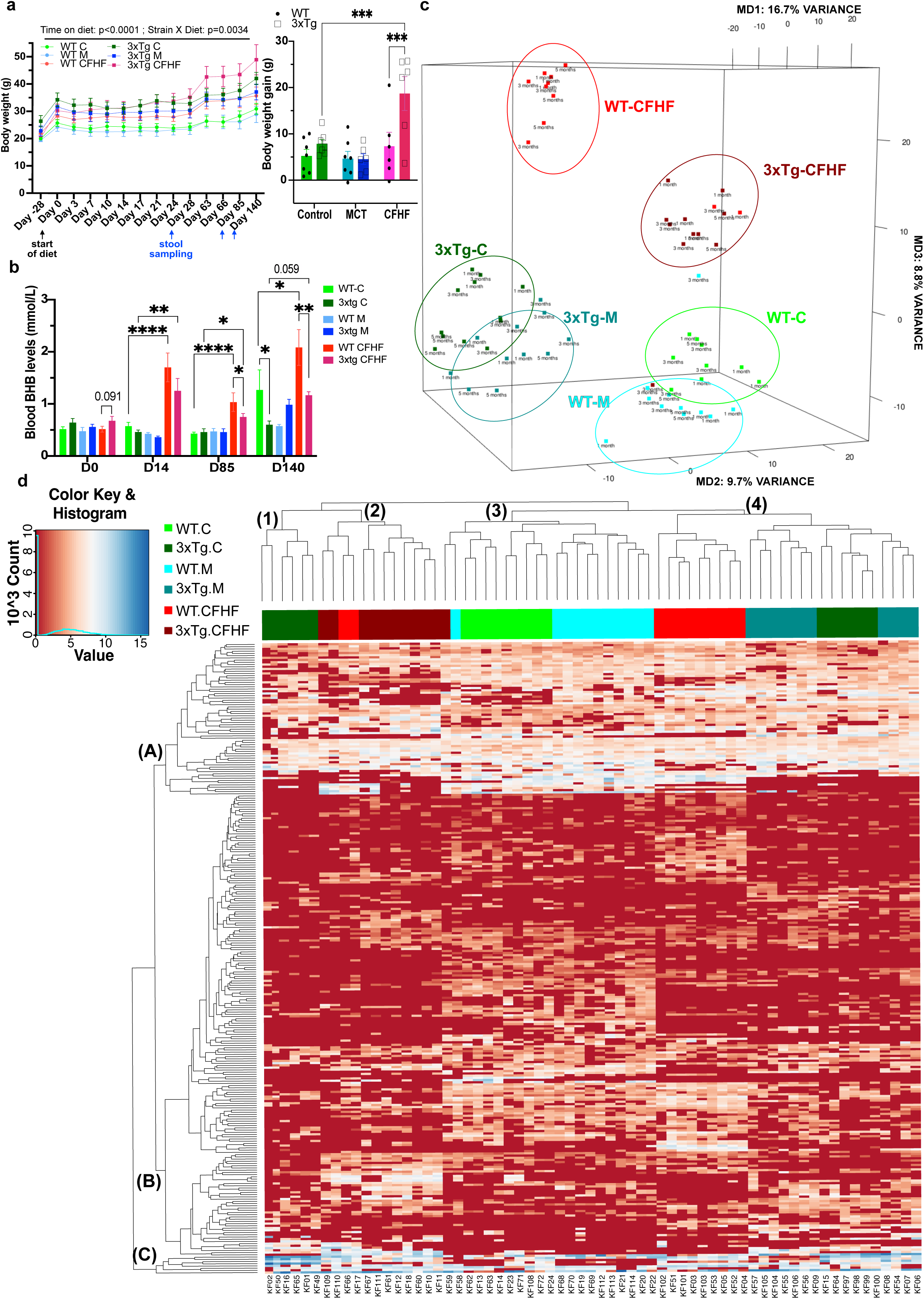
Monitoring of metabolic features and overview of faecal microbiome features in 3xTg-AD mice treated with ketogenic interventions. a) Body weight monitoring in mice fed diets (n=6-7/strain/diet). The mixed-effect stats model is based on following fixed effects: Time on diet (p-value<0.0001, F(2.473, 83.71) = 90.73), Strain X Diet (p-value=0.0034, F(5, 34) = 4.394), Interaction (Time x Strain X Diet, p-value<0.0001, F(65, 440) = 4.121). The arrows indicate the start of the diet (black) and the time when fecal matter was harvested (brown). b) Blood levels of beta-hydroxybutyrate(BHB) at different timepoints. The 2-way ANOVA is based on Strain and Diet factors: Day 0(D0): Strain (p-value = 0.0045, F(1, 36) = 9.203), Diet (p-value=n.s., F(2, 36) = 0.7099), Interaction(p-value=n.s., F(2, 36) = 1.318) Day14(D14): Strain(p-value =n.s., F(1, 34) = 2.028), Diet (p-value<0.0001, F(2, 34) = 31.76), Interaction(p-value=n.s., F(2, 34) = 1.469); Day85(D85): Strain (p-value =n.s., F(1, 34) = 1.688), Diet(p-value<0.0001, F(2, 34) = 17.04), Interaction(p-value=n.s., F(2, 34) = 1.903); Day104(D104) : Strain (p-value =0.017, F(1, 33) = 6.306), Diet (p-value=0.0009, F(2, 33) = 8.679), Interaction(p-value=0.0055, F(2, 33) = 6.127). c) Multidimensional scaling (MDS) analysis of microbiome of experimental groups (65 samples, n=3-4/strain/diet/time excepted WT-C-5 months = 1) with percentage of variance explained, legend indicate strain, diet and time d) Unbiased heatmap of all varying bacterial species abundance in faecal microbiome with hierarchical clustering of rows (283 ASVs) and columns (65 samples, n=3-4/strain/diet/timepoint excepted WT-C-5 months = 1), legend indicate strain, diet and level of abundance Significant pair-wise comparisons (uncorr. Fisher’s LSD) on graph: *p-value<0.05; **p-value<0.01; ***p-value<0.001; ****p-value<0.0001

We then analyzed the dissimilarities among the gut microbiomes using multidimensional scaling (MDS, **Fig.1c**, **Supp.Fig.1c**), hierarchical clustering, and a heatmap of the relative abundance of the most variable amplicon sequence variants (ASVs, **Fig.1d**, **Supp.Fig.1d**). MDS analysis showed a marked distinction between 3xTg-AD and WT mice samples, as visualized by their separation along the two main axes of the plot, which together captured 26.7% of the variability of the data (**Fig.1c**). The second main difference was among the dietary interventions. Mice fed the MCT and Control diets were slightly separated but strongly differed from mice fed the CFHF diet along the MD3 axis, explaining 8.8% of the variance (**Fig.1c**). Using the abundance patterns of 283 ASVs and an unsupervised clustering method, we classified 3xTg-AD and WT mice into four subclusters (**Fig.1d**). Based on the dendrogram, subclusters (1) and (2) are more similar to each other, whereas subclusters (3) and (4) are more similar to each other. Subcluster (1) comprised 3xTg-AD mice fed the Control diet at 4 and 6 months of age. Subcluster (2) mainly included 3xTg-AD mice of all ages fed a CFHF diet. Subcluster (3) consisted of WT mice fed the Control and MCT diets. Subcluster (4) regrouped WT mice on the CFHF diet, 3xTg-AD mice on the MCT diet, and 3xTg-AD mice on the Control diet mainly at 8 months of age (**Fig.1d**). The same analysis was performed for the 5xFAD model. The MDS plot showed a significant difference between CFHF-fed mice and other mice along the main axis (**Supp.Fig.1c**). Within CFHF-fed mice, there was a minor but clear distinction between 5xFAD mice and their WT counterparts (**Supp.Fig.1c**). The heatmap showed similar observations, where the abundance patterns of 218 ASVs allowed the identification of two clusters. One with the CFHF-fed mice and the second with the MCT and Control diets-fed mice (**Supp.Fig.1d**). Within cluster (1), we distinguished two subclusters separating WT mice on the CFHF diet from 5xFAD mice on the CFHF diet (**Supp.Fig.1d**).

These data reveal microbiome alterations linked to AD mutations, type of diet, and mouse age, the latter being observed only in 3xTg-AD mice.

### Assessments of alpha and beta diversity highlight diet effects on the microbiota of AD mice, especially at younger ages

Microbiome diversity is one of the first quantitative analyses of microbiome alterations. It can be assessed at different levels, with alpha and beta diversities being the primary diversities. Alpha diversity or within-sample diversity describes the number of different species (<u>richness)</u> and their relative abundance (<u>evenness</u> of species representation) within a specific sample ^39^. Beta diversity refers to diversity across samples, describing how similar or divergent there are two samples^39^. Here, we investigated the effects of ketogenic diets on microbiome alpha and beta diversities in transgenic AD mice treated at different time points.

Alpha diversity was analyzed using three indices, namely Shannon describing the evenness^39^, Chao1 for richness^39^, and Simpson evaluating the presence of dominant species over the others^40^. At 4 months of age, 3xTg-AD mice on the Control diet presented significantly lower Shannon and Chao1 indices than WT mice, indicating a less even distribution and reduced number of different species (**Fig.2a, 2c**). Both MCT and CFHF increased these indices in 3xTg-AD mice after 1 month of intervention without changing the values in WT mice (**Fig.2a, 2c**). The Simpson index did not reveal significant differences between WT and 3xTg-AD mice fed the Control diet (**Fig.2b**). Interestingly, 3xTg-AD mice on the CFHF diet presented a higher Simpson index than 3xTg-AD mice on the Control diet, indicating a higher predominance of specific species due to the CFHF diet (**Fig.2b**). At 6 months of age, Shannon and Simpson indices in 3xTg-AD mice did not differ from those in WT mice, while the Chao1 index remained lower (**Fig.2a-c**). At 8 months of age, there were no significant differences in these indices between WT and 3xTg-AD mice fed the Control diet (**Fig.2b**). Interestingly, after 3 and 5 months of intervention, MCT- and CFHF-fed mice did not differ from 3xTg-AD mice on the Control diet (**Fig.2a-c**). The alpha diversity indices of mice fed the MCT diet did not change with time (**Supp.Fig.3a-c**). 3xTg-AD mice on a CFHF diet for 1 month showed higher Shannon and Simpson indices than those treated for 3 months and were similar to those treated for 5 months (**Supp.Fig.3a-b**). Overall, dietary interventions with MCT and CFHF significantly elevated microbial alpha diversity in 3xTg-AD mice, particularly at younger ages, as evidenced by the increased Shannon and Chao1 indices, although these effects diminished with age and prolonged treatment duration.

**Figure 2.**
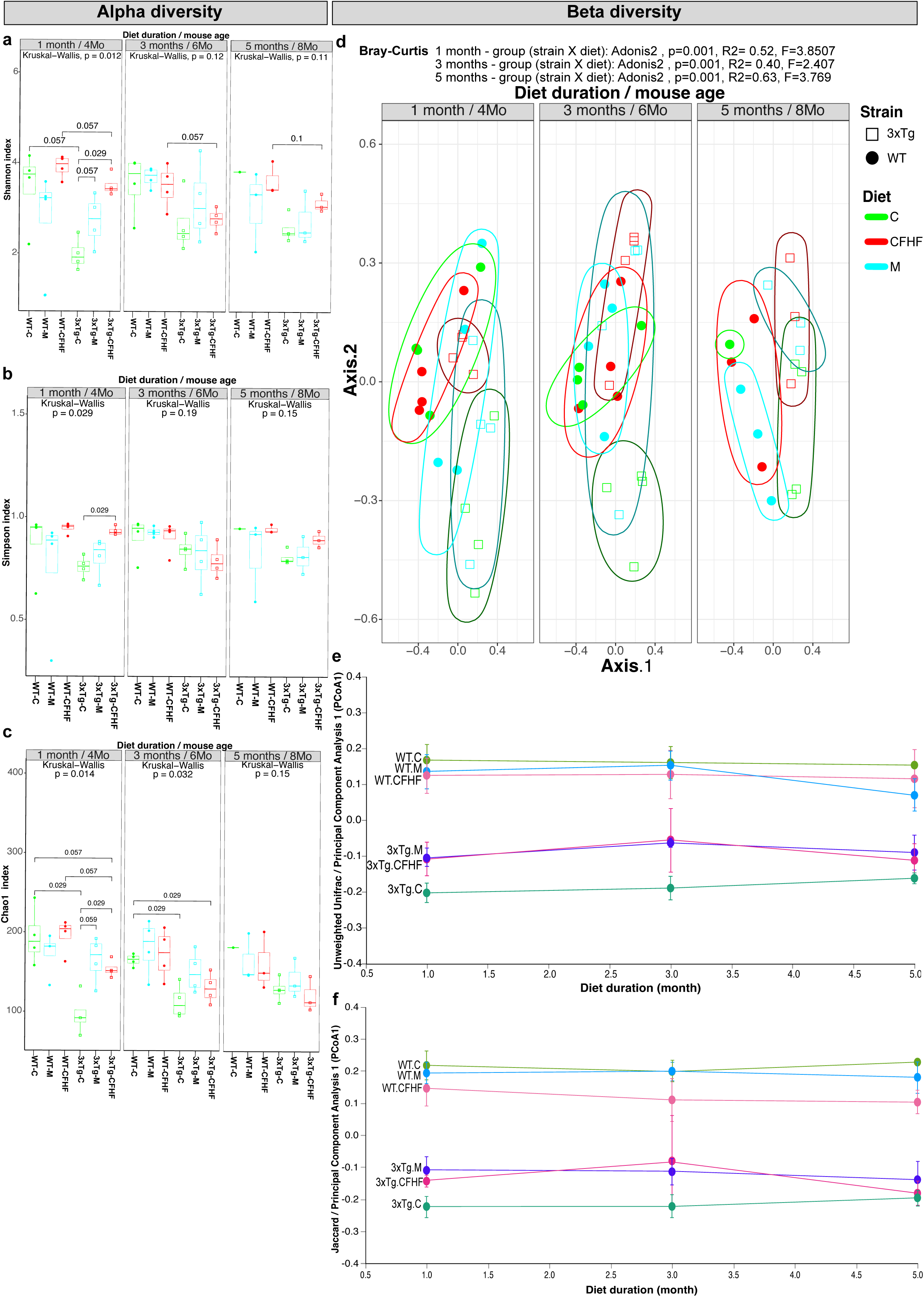
Longitudinal assessment of Alpha and Beta diversity of the microbiome in 3xTg-AD mice on standard diet and treated with ketogenic interventions. a-c)Indices of Alpha diversity at different time points of the diet including a)Shannon index, b)Simpson index, and c)Chao1 index (Multivariate statistical analysis with Kruskal Wallis and pair-wise comparison post-hoc test: Wilcoxon, resuts on graph). d-f) Indices of beta-diversity including d)Bray-Curtis showing PCoA1 and PCoA2 (Multivariate statistical analysis with Adonis2 considering the following factors : group(strainXdiet) and residual, results for group factor on graph), e) Volatility plot of the average changes of PCoA1 of Unweighted Unifrac distance matrix, f) Volatility plot of the average changes of PCoA1 of Jaccard distance matrix

**Figure 3.**
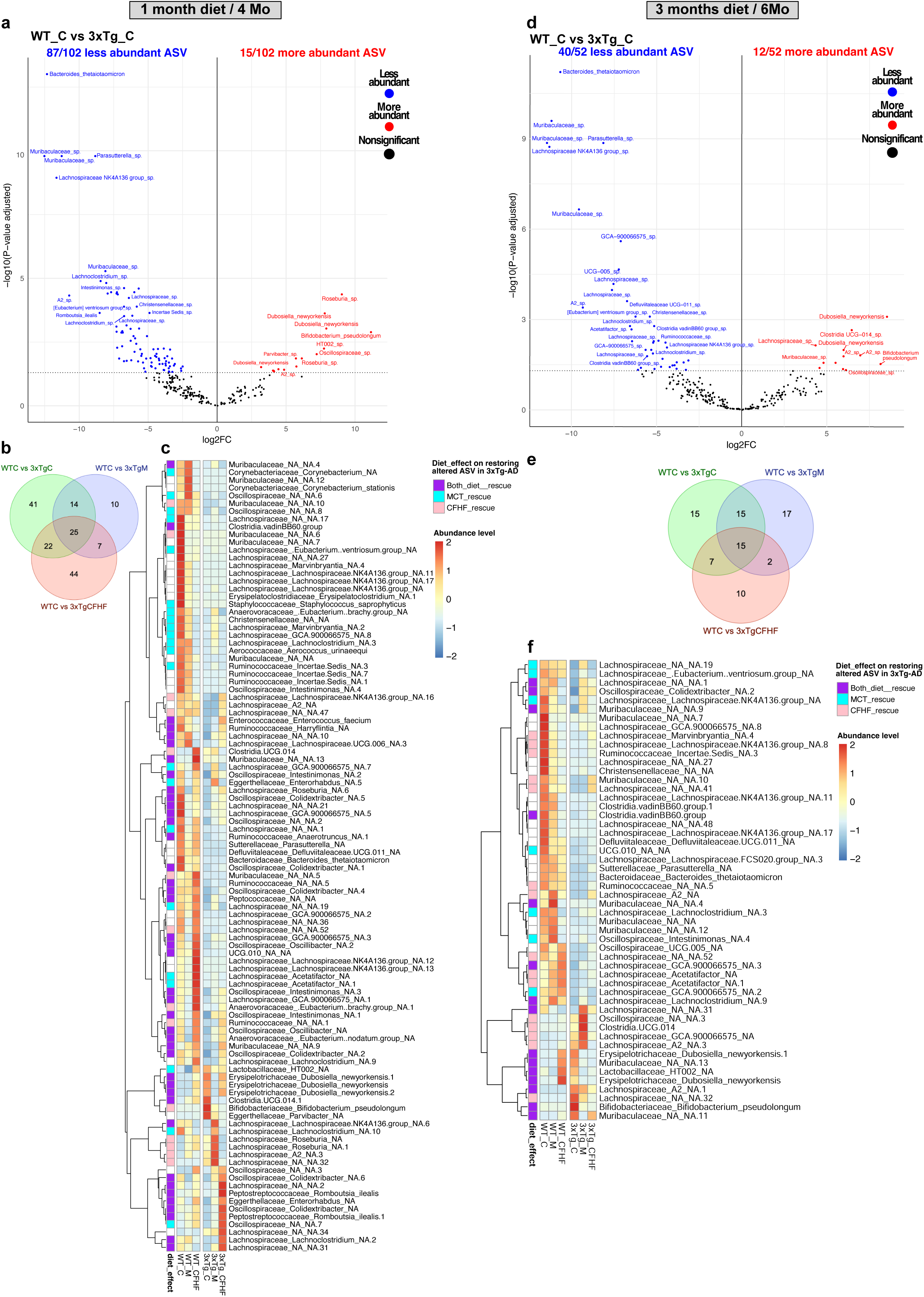
AD-induced altered relative abundance levels of several ASVs which are restored in part by both MCT and CFHF diets after 1 month and 3 months of intervention. **a-c)** Differential expression analysis after 1 month of ketogenic interventions, obtained with Limma (adjusted p-value <0.05) to **a)** visualize significant changes in relative abundance levels of ASVs between WT and 3xTg-AD mice on Control diet (WT C vs 3xTg C) with Volcano plot; **b)** evaluate the number of ASVs restored by MCT or CFHF diet by comparing the number of ASVs altered in the following contrasts in Venn diagram: AD-induced alterations (WT C vs 3xTg C), MCT effect in AD mice (WT C vs 3xTg M), CFHF effect in AD mice (WT C vs 3xTg CFHF). The ASVs that are significantly different in WT C vs 3xTg C and not significant in WT C vs 3xTg M or WT C vs 3xTg CFHF are considered “restored”; **c)** and visualize MCT and CFHF effects on the relative abundance of altered ASVs in WT C vs 3xTg C with a hierarchical heatmap. The color code in the extreme left column indicates whether the ASV is rescued by MCT (cyan), CFHF (pink), or both diets (purple). **d-e)** Differential expression analysis after 3 months of ketogenic interventions, obtained with Limma (adjusted p-value <0.05). Same graphs and corresponding explanation as in **Fig.3a-c**. (See Detailed raw numbers in Supp. Table 1)

We used three indices of beta-diversity: Bray-Curtis (based on relative abundance)^41^, Unweighted Unifrac (presence/absence of species and phylogenic information)^42^, and Jaccard (presence/absence of species)^41^. We report marked dissimilarity between WT and 3xTg-AD mice on the Control diet at each time point for all three indices (**Fig.2d-f**). Using the Bray-Curtis metric, we observed that the microbiome profile of 3xTg-AD mice on either the MCT or CFHF diet was closer to that of WT mice on the Control diet, especially after 1 month and 3 months of intervention (**Fig.2d**). Jaccard and unweighted Unifrac plots indicated that 3xTg-AD mice on CFHF and MCT diets resembled each other and differed from 3xTg-AD mice on Control diet. This difference tended to diminish with prolonged diet and age (**Fig.2e-f**). In contrast to the Bray-Curtis results, Jaccard and unweighted Unifrac analyses indicated that 3xTg-AD mice on CFHF and MCT remained distant from WT mice (**Fig.2e-f**). These results reveal that MCT and CFHF diets alter the richness of microbial species in 3xTg-AD mice to resemble that of WT mice but maintain a different species composition compared to WT mice. These effects were more significant in younger mice.

Alpha and beta diversity metrics of 5xFAD mice did not differ significantly from those of WT mice on the Control diet (**Supp.Fig.2**). However, 5xFAD and WT mice responded differently to ketogenic diets after one month of intervention. The MCT diet reduced alpha diversity indices in 5xFAD mice but not in WT mice, while CFHF reduced the Chao1 index in 5xFAD mice (**Supp.Fig.2a-c**). We did report significant differences in alpha diversity after 3 months of intervention (**Supp.Fig.2a-c**). All beta diversity metrics indicated a marked distinction between mice subjected to the CFHF diet and the other mice (**Supp.Fig.2d-f**). WT and 5xFAD mice on the CFHF diet showed dissimilarities with Bray-Curtis at 1 month of intervention and unweighted Unifrac at 3 months of intervention (**Supp.Fig.2d-e**). The unweighted UniFrac analysis also reported differences between WT and 5xFAD mice subjected to the MCT diet for 1 month (**Supp.Fig.2e**). In summary, although 5xFAD mice did not show microbial alterations at baseline, they exhibited specific microbial changes under ketogenic interventions.

Taken together, the microbiome diversity analysis revealed early alterations in alpha and beta diversities related to AD mutations. Ketogenic interventions partially prevented these changes.

### MCT and CFHF restored ASVs with differential abundance levels and induced specific alterations in AD mice

We investigated the bacterial species associated with AD mutations and ketogenic interventions by examining the levels of bacterial abundance. We performed this analysis at the ASV, phylum, and family levels.

Of the 283 variable ASVs, differential expression analysis revealed that 102 ASVs were significantly different between WT mice and 3xTg-AD mice on the Control diet at 4 months of age (**Fig.3a**). 87/102 were less abundant in 3xTg-AD mice, whereas 15/102 were more abundant (**Fig.3a**). The most abundant was *Bifidobacterium pseudolongum,* tenfold higher in 3xTg-AD mice. The least abundant were *Bacteroides thetaiotaomicron* and another species from the family *Muribaculaceae*, each of them from the *Bacteroidetes phylum*, and twelve-fold less abundant in 3xTg-AD mice (**Fig.3a**). At phylum level, we observed that 3xTg-AD mice exhibited higher levels of *Actinobacteriota* owing to the abundance of *Bifidobacterium pseudolongum* from *the Bacteroidaceae* family (**Fig.4a, 4c**). *Bacteroidota* and *Proteobacteria* were globally in low abundance in 3xTg-AD mice and most of them were not restored by diet (**Fig.4a**). 3xTg-AD mice did not present alterations in overall level of *Verrumicrobiota* and *Firmicutes*(**Fig.4a**). 3xTg-AD mice presented higher *Firmicute:Bactoroidota* ratio (F/B), indicator of an altered microbiome (**Fig.4a**). Finally, we studied the families constituting *Firmicutes*, which showed alterations in *Lachnospiraceae*, *Erysipelotrichaceae*, *Lactobacillaceae*, *Oscillospiraceae*, *Oscillospirales type UCG-010*, and *Anaerovoracaceae* in 3xTg-AD mice (**Supp.Fig.6, Fig.3c**). Three *Firmicutes* with unidentified families were also altered in 3xTg-AD mice and belonged to the orders *Clostridia UCG-014* and *Clostridia vadin BB60* (**Fig.3c**).

**Figure 4.**
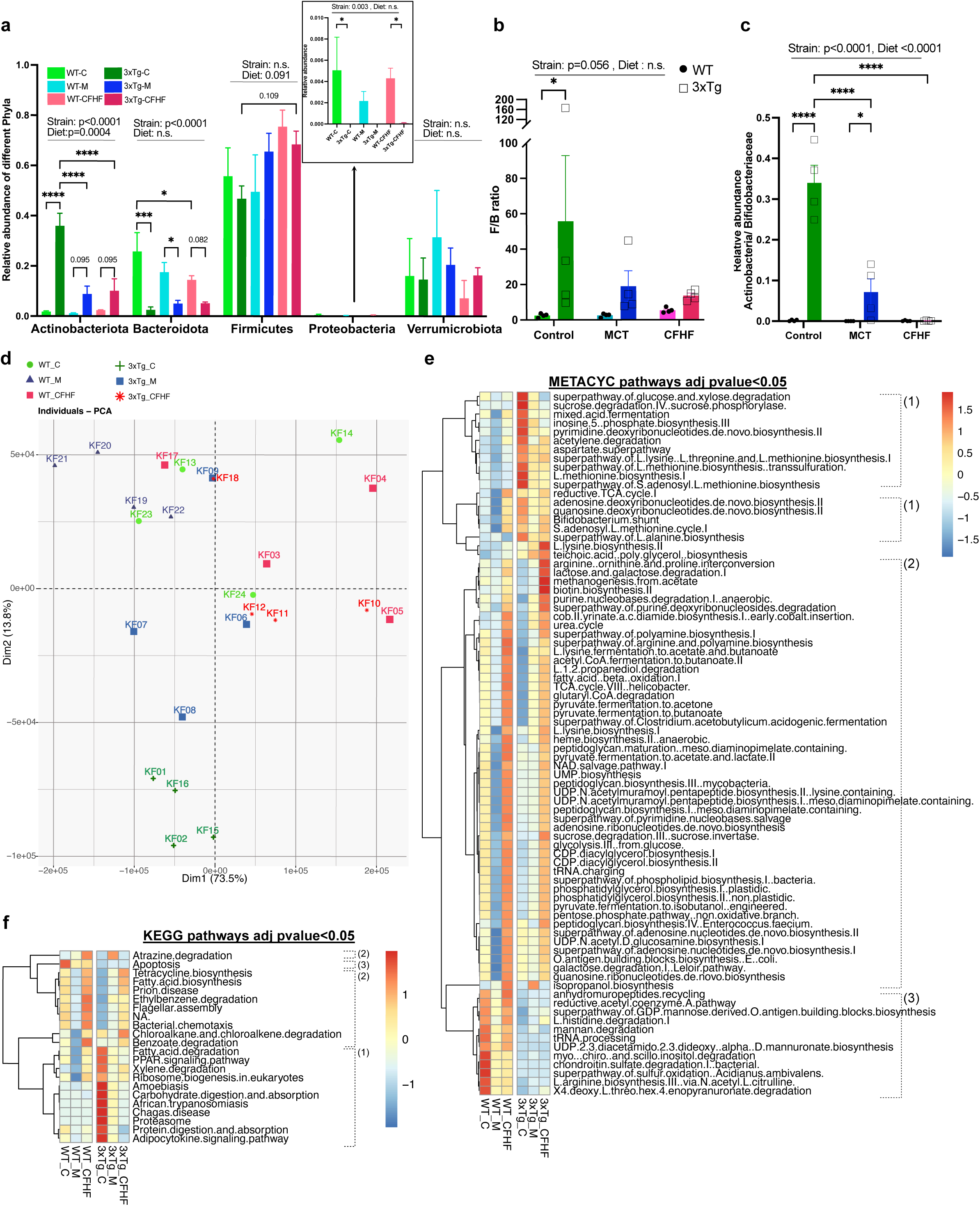
Microbial alterations at phylum and family levels and PICRUST predicted functional changes in the microbiome based on 16S rRNA sequencing after 1 month of ketogenic interventions. **a-c)** Relative abundance levels of bacteria at phylum and family levels after 1 month of ketogenic interventions showing **a)** identified phyla alterations (Stats on graph : For each phylum - 2way-ANOVA based on Strain and Diet factors and Uncorrected Fisher’s LSD for pair-wise comparison tests), **b)** Ratio of *Firmicutes:Bacteroidetes* (F/B ratio, Stats on graph : 2way-ANOVA based on Strain and Diet factors and Uncorrected Fisher’s LSD for pair-wise comparison tests), **c)** Relative abundance of the most abundant family in 3xTg-AD mice and diet effect : *Bifidobacteriaceae* (Stats on graph : 2way-ANOVA based on Strain and Diet factors and Uncorrected Fisher’s LSD for pair-wise comparison tests). **d-e)** PICRUST analysis annotated with Metacyc database showing **d)** Principal Component Analysis (PCA) of each sample (labelled KF01 to KF24) and e) Heatmap of significant pathways with differential abundance (ALDEx2 Kruskal-Wallace, adjusted p-value <0.05). **f)** PICRUST analysis annotated with KEGG database showing heatmap of significant pathways with differential abundance (ALDEx2 Kruskal-Wallace, adjusted p-value <0.05) The pathways were classified into 3 groups based on their abundance: (1) pathways enriched in 3xTg-AD mice compared to WT mice on Control diet and reduced by MCT or CFHF diet, (2) pathways less enriched or unaltered in 3xTg-AD mice compared to WT mice on Control diet and increased by MCT or CFHF and finally (3) pathways altered in 3xTg-AD mice with no diet effect. Star legend: *p-value<0.05; **p-value<0.01; ***p-value<0.001; ****p-value<0.0001

Administration of the MCT diet for 1 month corrected the relative abundance of 63/102 ASVs (**Fig.3b**). MCT partially restored the levels of *Bifidobacterium pseudolongum* (**Fig.3c**, **Fig.4a, 4c**) as well as several types of *Firmicutes* (**Fig.3c**, **Supp.Fig.6**). MCT also altered the relative abundance of 17 ASVs that were not altered in 3xTg-AD mice on the Control diet (**Fig.3b**). CFHF corrected 55/102 ASVs (**Fig.3b-c**) including total restoration of the *Bifidobacterium pseudolongum* levels toward WT mice level (**Fig.3c**, **Fig.4a,4c**) and overall levels of *Lachnospiracea*e, *Erysipelotrichaceae*, and *Lactobacillaceae* (**Supp.Fig.6**). CFHF induced changes in the relative abundance of 51 new ASVs (**Fig.3b**).

At 6 months of age, 52 ASVs were altered in 3xTg-AD mice on the Control diet, with 40 being less abundant and 12 being more abundant compared to the WT mice microbiome (**Fig.3d**). The species types affected resembled those seen in 4 months old 3xTg-AD mice, with 83% (43/52) altered ASVs in common (**Supp.Fig.6**). Both ketogenic interventions until this age restored 15 ASVs, with MCT restoring the level of seven more (22 ASVs restored in total, **Fig.3e**), and CFHF restoring 15 more (30 ASVs restored in total, **Fig.3e**). *Bifidobacterium pseudolongum* was completely restored by both interventions (**Fig.3e**), while *Bacteroides thetaiotaomicron* remained unchanged by diet (**Fig.3e**).

As the microbiome diversity of 5xFAD mice appeared similar to that of WT mice on the Control diet, we found only eight ASVs that were altered in 5xFAD mice at 3 months of age (**Supp.Fig.4a-b**) and only one at 5 months of age. At the phylum level, 5xFAD mice at 3 months of age had higher levels of *Proteobacteria* associated with *Enterobacteriaceae*, such as *Escherichia coli (shigella)* (**Supp.Fig.4a, Supp.Fig. 5**), and *Moraxellaceae acinetobacter schindleri* (**Supp.Fig. 5**), which was not among the most variable ASVs (**Supp.Fig.1d**). CFHF restored all altered ASVs in 3 months old 5xFAD mice, while MCT restored 6/8 ASVs (**Supp.Fig.4b**). The most abundant ASV was a *Muribaculum* species, which was also the only one altered in 5 months old 5xFAD mice. It was restored by both diets during both periods (**Supp.Fig.4a-b**). Interestingly, 5xFAD mice presented specific features when exposed to the MCT or CFHF diet, especially *Firmicutes* and *Verrumicrobiota* (**Supp.Fig.5a**). After 1 month of intervention, MCT tended to decrease the levels of *Firmicutes* more in 5xFAD mice than WT mice (**Supp. Fig.5a**), primarily affecting *Lachnospiraceae* and *Oscillospiraceae* (**Supp.Fig.4c, Supp. Fig.5c**). The same trend was observed at 3 months post-intervention, yet with less ASVs in both WT and 5xFAD mice (**Supp.Fig.4e**). CFHF increased *Firmicutes* and decreased *Verrumicrobiota* in 5xFAD mice, while doing the opposite in WT mice (**Supp. Fig.5a, Supp. Fig.4d**). This resulted in a significant increase in *F/B* in 5xFAD mice fed the CFHF diet compared to WT mice fed the same diet (**Supp. Fig.5b**). *Akkermansia municiphila* is the most common *Verrumicrobiota* in the microbiome, which was significantly reduced in 5xFAD mice fed a CFHF diet (**Supp.Fig.4d**). The same differential response was observed at 3 months post-intervention, but with reduced ASVs (**Supp.Fig.4f**).

**Figure 5.**
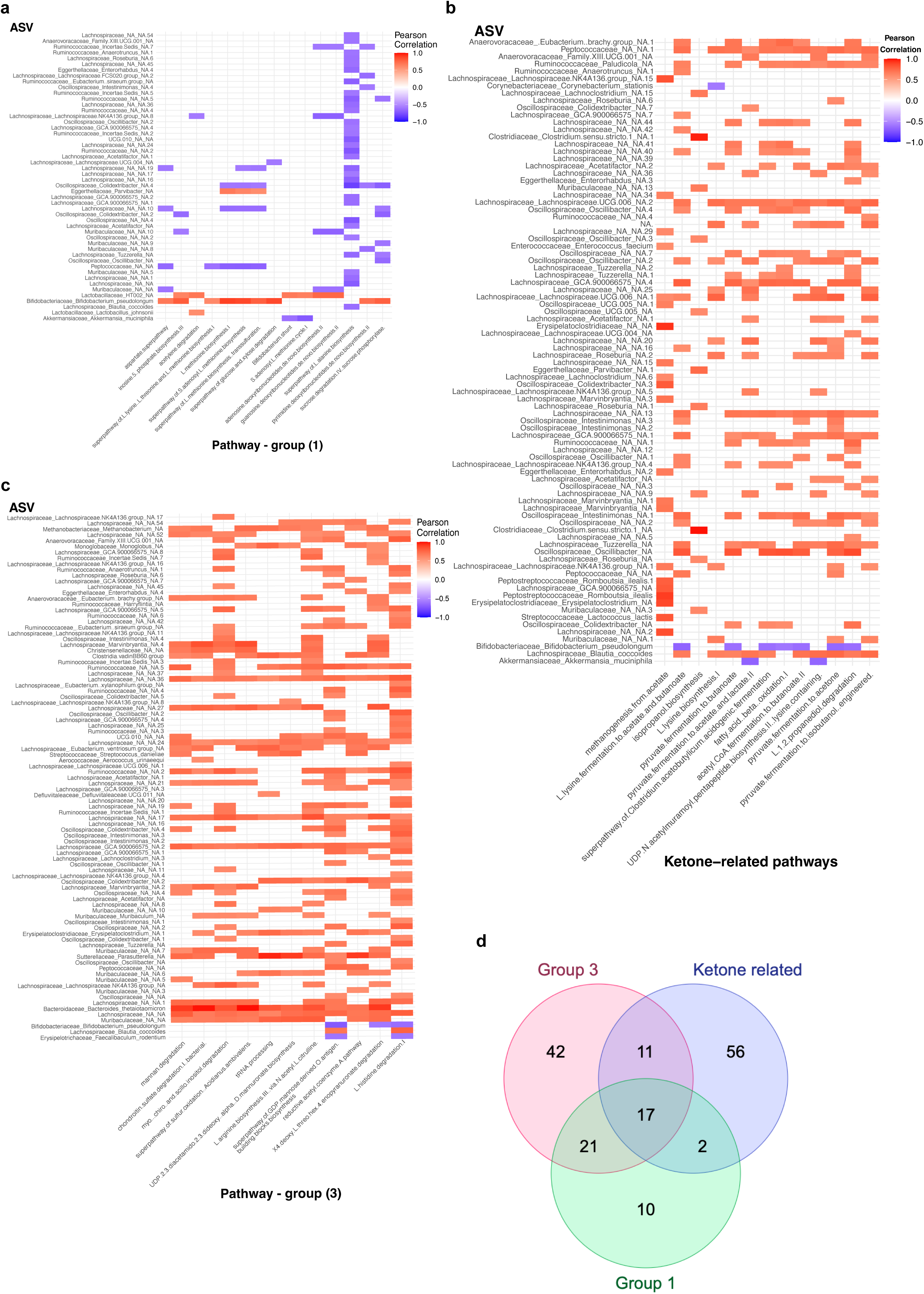
Correlation between varying ASVs and predicted functional pathways. **a-c)** Heatmap showing the correlation between ASVs and predicted functional pathways classified into 3 groups : **a)** Pathways from the group(1) enriched in 3xTg-AD mice compared to WT mice on Control diet and reduced by MCT or CFHF diet, **b)** pathways from the group (2) : less enriched or unaltered in 3xTg-AD mice compared to WT mice on Control diet and increased by MCT or CFHF. Here, we show the ones that are directly associated with ketone or SCFA metabolism pathways, and **c)** pathways from group (3) altered in 3xTg-AD mice with no dietary effect. (See raw numbers in Supp. Table 3) **d)** Venn diagram of ASVs correlating with each pathway group to evaluate dissimilarities.

This shows that genetic models of AD present alterations in various types of bacteria, including *Actinobacteria, Bacteroidetes, Firmicutes, and Verrumicrobiota.* Ketogenic interventions shifted the levels of these bacteria in AD mice to normal conditions (WT mice fed the Control diet). However, mice subjected to MCT or CFHF diets presented other alterations associated with diet effects and their genetic background.

### Functional mechanisms associated with changes in the microbiome of 3xTg-AD mice fed with MCT or CFHF diet for 1 month

Different microbiome diversities do not always imply functional changes, as different bacteria may have the same functions. Here, we used PICRUST to predict functional changes associated with the microbiome composition of 3xTg-AD mice fed a Control diet and treated with ketogenic interventions for 1 month. We used two databases, Metacyc and KEGG, to report metabolic pathway profiles.

Using the metabolic pathway profiles obtained with Metacyc, we performed principal component analysis (PCA) showing a distinction between WT and 3xTg-AD mice on the Control diet along axis 2 (Dim 2-bottom/up, **Fig.4d**). The 3xTg-AD mice on MCT or CFHF diets were closer to the WT mice fed the Control diet along the same axis (Dim 2, **Fig.4d**). Most mice fed the CFHF diet were separated from those fed the MCT diet on axis 1, but with some variability (Dim1-right/left, **Fig.4d**). This indicates differential pathways according to genetics and diet.

80 pathways identified with Metacyc and 21 pathways identified with KEGG were significantly different among the six experimental groups (**Fig.4e-f**). These pathways can be divided into three main groups: (1) pathways enriched in 3xTg-AD mice compared to WT mice on the Control diet and reduced by MCT or CFHF diet, (2) pathways that were less enriched or unaltered in 3xTg-AD mice compared to WT mice on the Control diet and increased by MCT or CFHF, and (3) pathways altered in 3xTg-AD mice with no diet effect (**Fig.4e**). For the first group, Metacyc and KEGG analyses revealed 26 pathways that were enriched in 3xTg-AD mice on the Control diet and reduced in 3xTg-AD mice on the MCT or CFHF diet. These pathways were related to <u>amino acids</u> (protein digestion and absorption, synthesis of aspartate-derived amino acids (L-lysine, L-threonine, L-methionine), L-alanine biosynthesis, and proteasome), <u>carbohydrates (</u> carbohydrate digestion and absorption, sucrose and glucose degradation), and <u>synthesis of DNA precursors</u> (pyrimidine and inosine monophosphate biosynthesis) (**Fig 4e-f, supp.Fig.6c**). The precursors of all these pathways are produced in the reductive tricarboxylic acid (TCA) pathway^43,44^, which was also enriched in 3xTg-AD mice but was unaltered by ketogenic diets (**Fig.4e**). KEGG revealed increased lipid mechanisms, including PPAR signaling, fatty acid degradation, adipocytokine signaling, and Metacyc, highlighted increased *Bifidobacterium shunt* (**Fig.4e**) and lactate production from hexose, also occurring in *Bifidobacteria*^44,45^.

In the second group, both databases highlighted a decrease in lipid biosynthesis in the microbiome of 3xTg-AD mice fed the Control diet, notably fatty acid biosynthesis, phospholipid biosynthesis, diacylglycerol, and phosphatidylglycerol. These pathways were upregulated by the CFHF diet (**Fig.4e-f**). 3xTg-AD mice fed MCT and/or CFHF diets had a microbiome enriched with ketone bodies or short-chain fatty acid (SCFA)-related pathways, including butanoate and acetoacetate production from pyruvate, acetyl-CoA, and L-lysine fermentation, as well as propionate formation from propane-diol degradation and isopropanol biosynthesis (**Fig.4e**). These pathways related to ketone body substrates or SCFA substrates, such as L-lysine biosynthesis and fatty acid beta-oxidation (**Fig.4e**), as well as related bacteria, such as *Clostridium acetobutylicum* (**Fig.4e**). Both diets resulted in an increased urea cycle and purine degradation (**Fig.4e**). CFHF, but not MCT, increased various pathways of glycolysis and sucrose degradation and several arginine- and polyamine-related pathways (**Fig.4e**).

Finally, approximately 12 pathways were less enriched in 3xTg-AD mice on the Control diet and remained unaffected by ketogenic diets, with the most significant being apoptosis (**Fig.4f**), mannan degradation, chondroitin degradation, and inositol degradation (**Supp.Fig.6**, **Fig.4e**).

These data reveal functional changes associated with the microbiome of 3xTg-AD mice. These changes involve macronutrient and nucleotide metabolism that were regulated by the CFHF and MCT diets. Ketogenic interventions stimulated SCFA production and upstream pathways.

### Bacterial changes correlate with predicted functional mechanisms altered by AD genetic and ketogenic interventions

To identify which bacteria might be involved in the functional changes predicted in 3xTg-AD mice, we investigated the correlation between variable ASVs and potentially altered pathways. Using the previous pathway classification, 50 ASVs were significantly associated with group (1) (increased in 3xTg-AD mice on Control diet and reduced by MCT or CFHF, **Fig.5a**), 91 ASVs with group (3) (decreased in 3xTg-AD mice on Control diet with no diet effect, **Fig.5c**), and 115 ASVs with group (2) (decreased or not affected in 3xTg-AD mice on Control diet and stimulated by MCT or CFHF, **Fig.5b**, **Supp.Fig.8).** This group was subdivided into two groups: ketone-related pathways (**Fig.5b**) and non-ketone pathways (**Supp.Fig.8**). Pathways that were not significantly associated with any ASV were removed (**Supp. Table 3**).

Interestingly, the *Bifidobacterium pseudolongum,* which was the most abundant ASV in 3xTg-AD mice on the Control diet and was significantly reduced by both ketogenic interventions (**Fig.3a**, **Fig.3c**, **Fig.4a**), was positively correlated with 9/15 pathways increased in 3xTg-AD mice on the Control diet (**group (1), Fig.5a**), negatively correlated with 3/11 pathways always decreased in 3xTg-AD mice (**group (3), Fig.5c**), and also negatively correlated with 7/13 ketone-related pathways (**group (2), Fig.5b**). Similarly, the least abundant ASV in 3xTg-AD mice, independently of diet group named *Bacteroides thetaiotaomicron* (**Fig.3c**) and a species of *Muribaculaceae (Muribaculaceae sp.,* ASV10, **Supp. Table 1, Fig.3c**) were positively correlated with almost all pathways that were less enriched in 3xTg-AD mice (**group(3), Fig.5c**). The same Muribaculaceae species was also negatively correlated with pathways enriched in 3xTg-AD mice (*Muribaculaceae sp.,* **group (1), Fig.5a**).

By analyzing the pathways involved in group (1), we noted that the mechanisms of methionine biosynthesis were positively correlated with a species of *Eggerthellaceae Parvibacter sp.* (**Fig.5a**). The *Bifidobacterium* shunt and the biosynthesis of DNA precursors were associated with a species of *Lactobacillaceae HT002 sp.* (**Fig.5a**). *Lactobacillus johnsonii* was positively correlated with acetylene degradation (**Fig.5a**). This is in line with increased levels of *Lactobacillaceae* in 3xTg-AD mice on the Control diet and reduced levels of this family in 3xTg-AD mice under ketogenic interventions (**Supp.Fig.6**).

46% (42/91) of the ASVs correlating with pathways in group (3) were *Lachnospiraceae*, with some of them being positively correlated with all these pathways *(Lachnospiraceae sp. ; Lachnospiraceae sp.36; Lachnospiraceae sp.17; Lachnospiraceae sp.27; Lachnospiraceae GCA.S000CC575 sp. …* **Supp. Table 1**, **Fig.5c**). This is in line with the reduced levels of *Lachnospiraceae* observed in 3xTg-AD mice on the Control diet (**Supp. Fig.6**). *Oscillospiraceae* represented the second most affected ASVs in group (3) with 15% (14/91) of them being positively correlated with the concerned pathways (**Fig.5c**).

In group (2), the ketone-related pathways were positively correlated mainly with species of *Lachnospiraceae* and *Oscillospiraceae.* Most of these species were not correlated with pathways in group (1) or group(3) (**Fig.5d**), suggesting the specificity of bacteria and functional changes induced by ketogenic interventions. Five ASVs were positively correlated with almost all ketone-related pathways (>9/13). Those are : *Peptococcaceae sp.1, Ruminococcaceae Paludicola sp., Lachnospiraceae Lachnospiraceae.UCG.006 sp.2, Lachnospiraceae sp.13, Lachnospiraceae Blautia coccoide* (**Fig.5b**). The same ASVs were also positively correlated with almost all other pathways enriched by ketogenic interventions (**Supp. Fig. 8**).

Interestingly, isopropanol biosynthesis was highly correlated with two species of *Clostridium sensu stricto1* (**Fig.5b**, **Supp. Table 3** *Clostridium sensu stricto1 sp.* Pearson correlation = 0.99, *Clostridium sensu stricto1 sp.1* Pearson correlation = 0.96). This kind of bacteria and associated pathways were specifically enriched in 3xTg-AD mice on the MCT diet (**Fig.4e**, **Supp. Table 1**).

This section highlights the bacteria correlating with functional changes induced by AD genetic mutations and ketogenic interventions. This provides an idea of bacteria having potential causal effects on these mechanisms and how these bacteria would change the microbiome in individuals with AD treated with ketogenic interventions.

## DISCUSSION

This study reports the effects of ketogenic interventions on the fecal microbiome in transgenic mouse models of AD.

### MCT and CFHF prevent early alterations of the microbiome and induce a shift toward SCFA-pathways

1 month of MCT or CFHF intervention was sufficient to restore, respectively, 62% and 54% of disturbed ASVs in the microbiome of 3xTg-AD mice. These effects persisted after 3 months of treatment. In 5xFAD mice, MCT and CFHF induced specific alterations in their microbiome.

Both MCT and CFHF increased the alpha diversity of the microbiome in 3xTg-AD mice, whereas they either reduced or did not alter this parameter in 5xFAD mice. Both ketogenic interventions reduced the levels of *Bifidobacterium pseudolongum* in 3xTg-AD mice, and only CFHF depleted their abundance in 5xFAD mice microbiome. The abundance of *Bifidobacterium pseudolongum* was negatively correlated with ketone bodies and SCFA-related pathways. Although the link between *Bifidobacteria* and cognition is not clearly established, ketogenic interventions seem to reduce their levels^46^ potentially through the production of ketone bodies. Studies reporting the anti-seizure effects of ketogenic interventions in epilepsy also reported decreased levels of *Bifidobacteria* after a high-fat low-carbohydrate ketogenic diet ^17^. A small clinical trial evaluating the effect of a low-carbohydrate modified-Mediterranean ketogenic diet in MCI patients noticed a reduction in *Bifidobacteria^3C,^*^47^ in their patients, and similar observations were made in obese men treated with a low-carbohydrate ketogenic diet^48^. In this last study, it was proven that BHB directly inhibits the growth of different species of *Bifidobacteria*, which results in the decrease of pro-inflammatory immune cells in the gut and preserves the intestinal barrier^48^. Our study is the first to report such potential inhibitory effect of MCT supplementation on *Bifidobacteria*.

We observed that both diets also decreased the levels of *Lactobacillaceae*, especially *Lactobacillaceae HT002* and *Lactobacillaceae Johsonii,* in 5xFAD and 3xTg-AD mice. This has been reported in other studies in humans and mice treated with low-carbohydrate ketogenic diets^48–50^. MCT and CFHF reduced the levels of *Escherichia coli (shigella)* in 5xFAD mice, whereas several studies reported the opposite^17,48^. Ketogenic interventions also increased the levels of several SCFA-producing bacteria, such as *Lachnospiraceae* ^51^ and *Muribaculaceae*^52^, in 3xTg-AD and 5xFAD mice. Further investigations are needed to understand the downstream consequences of such changes, especially in AD.

### Different microbial alterations depending on the type of ketogenic intervention : MCT vs CFHF

The MCT diet is an alternative ketogenic intervention using MCFA to boost ketone production ^10,12,53^ and thus avoid the energetic consequences of carbohydrate deprivation. It is interesting to observe how a small proportion of MCFA (10%) incorporated into a high-carbohydrate diet can produce similar effects on the microbiome as a low-carbohydrate, high-fat diet-like intervention (CFHF). However, the two interventions differed. First, mice subjected to the MCT diet did not show a permanent plasma increase in BHB. The increase in SCFA-producing bacteria can be due to the presence of transient mild ketosis, as discussed in other studies^53–55^. Second, we observed differences in the bacterial and SCFA-related pathways driven by the MCT diet compared to CFHF. The 3xTg-AD and 5xFAD mice on the MCT diet had reduced global levels of *Lachnospiraceae* compared to those on the CFHF diet. Furthermore, in 3xTg-AD mice, MCT increased the levels of specific genera of *Lachnospiraceae* including *Roseburia, Lachnoclostridium, Lachnospiraceae GCA.S000CC575, Marvinbryantia,* and the *Clostridium* genus *Clostridium sensu stricto1.* Some of these genera, especially *Clostridium sensu stricto*, were correlated with isopropanol biosynthesis predicted to be increased in 3xTg-AD mice on an MCT diet according to PICRUST analysis. There has been no existing study on the microbiome of AD patients treated with MCT diets. However, studies reported that high-dose of MCT supplementation in mice on a standard diet resulted in increased levels of *Lachnospiraceae^5C^.* A high-fat MCT, low-carbohydrate diet in a Parkinson’s disease mouse model increased some species of *Lachnospiraceae* and *Clostridia*^57^ with no indication about the genus.

*Clostridium* species catalyze isopropanol biosynthesis from pyruvate, which is first metabolized into acetone and then isopropanol^44,58^. This suggests that the MCT diet promotes the production of isopropanol instead of the production of lactate from pyruvate, which would have been observed in a normal carbohydrate-enriched diet. Further investigations are required to understand the function of this reaction and the benefits in the context of AD.

### Genetic mutations of AD neuropathologies trigger alterations of bacteria relative abundances at younger ages

We have shown that 4 months old 3xTg-AD mice on a standard diet presented alterations in the microbiome, which tended to decrease with aging. These mice had lower microbiome diversity, a distinct microbiome composition compared to age-matched WT mice, and altered relative abundance of 102 ASVs at 4 months of age. Among these ASVs, we have reported increased levels of *Bifidobacterium pseudolongum* and reduced levels of *Bacteroidota,* resulting in a higher F/B ratio, and altered abundance in several families of *Firmicutes*. In 3 months old 5xFAD mice, we observed minor differences compared to WT mice, which mainly disappeared at 5 months of age. Young 5xFAD mice microbiome was characterized by enriched levels of *Esherichia coli (shigella), Muribaculaceae, and Enterococcaceae Enterococcus*, as well as decreased levels of *Defluviitaleaceae UCG 011 sp., Peptostreptococcaceae Romboutsia ilealis.1, Eggerthellaceae Parvibacter,* and altered levels of two species of *Lachnospiraceae*.

Similar to our observations, early changes in the microbiome of 3xTg-AD mice were shown to resorb over time in a longitudinal study from 4 to 13 months of age^23^. Changes in the gut microbiome in people at a high risk of developing AD dementia have been analyzed by few studies. Cognitively normal individuals with cerebral amyloid beta deposition presented minor alterations and no differences in alpha or beta diversity compared with those with no amyloid deposition^59^, similar to what was observed in our 5xFAD mice. Other studies reported changes in individuals with mild cognitive impairment (MCI) such as lower levels of *Bacteroidetes*^20,*C0,C1*^, higher levels of *Esherichia^C0^, Enterobacteriaceae*^20^*, Bifidobacterium^C0^*, *Lachnospiraceae Roseburia^C0,C1^*, and *Lactobacillus(Lactobacillaceae)^C0^,* as observed in the present study. A lower alpha diversity associated with reduced *Firmicutes* and increased *Bacteroidetes* was also reported in individuals with subjective cognitive decline and MCI^62^. In symptomatic AD individuals, the existing studies reported lower alpha-diversity and clear distinct microbiome alterations with the most consistent changes being reduced *Firmicutes^1S,C3,C4^* especially *Lachnospiraceae*^32^*^,C5-C7^* and *Clostridiaceae* families, reduced *Bacteroidota* ^22^*^,C3,C5,C8^* and increased *Escherichia coli*^22^*^,C3,C4^*. Interestingly *Bifidobacterium* levels were reported either increased or decreased in AD or MCI patients^18,19,21,32,60,65,67^. Strains of *Bifidobacterium* are often considered beneficial for gut health and are used in probiotic cocktails^45^. Here, we highlight the fact that these bacteria are increased by genetic risk factors for AD; therefore, more research is needed to understand their role in AD.

### AD mutation-induced alterations of bacteria levels are associated with potential functional changes in the microbiome

The predicted functional changes based on alterations in the microbiome of 3xTg-AD mice suggest impaired metabolic processes of macronutrients. *Bifidobacterium pseudolongum* and *Lactobacillaceae HT002* were associated with increased L-aspartate, L-threonine, L-lysine, and L-methionine biosynthetic pathways, sucrose degradation, hexose and xylose fermentation to pyruvate/lactate, and deoxyribonucleotide synthesis. Reduced levels of *Bacteroidota*, especially *Bacteroides thetaiotaomicron and Muribaculaceae,* correlated with a decrease in the degradation of mannan, inositol, chondroitin sulfate, and 4-deoxy-L-threo-hex-4-enopyranuronate. Some of these correlations have been proven in the literature^45,69^, for instance, the degrading effect of *Bacteroides thetaiotaomicron* on chondroitin sulfate^69^.

Some altered pathways observed in this study have been described in the brain, body or gut microbiome in the context of AD. For instance, higher microbial abundance of the sucrose degradation(sucrose phosphorylase) and s-adenosyl-methionine (SAMe) cycle 1 pathways has also been reported in MCI patients when compared to demented ones^68^. The 4-deoxy-L-threo-hex-4-enopyranuronate degradation^44^ another sugar catabolic pathway downstream of chondroitin sulfate degradation^44^and implicated in SCFA production^69,70^ was also disturbed in AD and MCI^64,71^. Chondroitin sulfate derived proteoglycans play a role in the response to neurological damages and might be involved in AD^72^. Nevertheless, nothing is known about the microbial chondroitin sulfate. Methionine levels and molecules involved in its cycle, such as homocysteine, have also been documented in patients with AD and MCI, as well as in interventions targeting this compound^73–75^. For instance, a genetic model of methionine restriction in drosophila highlighted the importance of microbiota-derived methionine in aging^75^.

In addition, our results also support the idea that the AD microbiome is characterized by a shift from SCFA-producing bacteria (*Bacteroidetes*, *Lachnospiraceae*) toward lactate-producing bacteria (*Bifidobacterium, Lactobacillaceae*)^32,67,76,77^. Reduced fecal levels of SCFA along different stages of AD^78,79^ have been reported. Notably, a study in APP/PS1 mice described excessive levels of lactic acid and reduced levels of SCFAs in their fecal matter^80^. Altogether, the onset of microbial alterations during early or pre-symptomatic stages of AD suggests a contribution of microbial species to AD pathophysiology through alterations in sugar, lactate, SCFA, and amino acids.

### Genetic mutations of AD influence their response to ketogenic interventions

Both 3xTg-AD and 5xFAD mice showed reduced upregulation of BHB and higher body weight gain under the CFHF diet compared to WT mice on the same diet. Furthermore, the microbiome composition of 5xFAD mice fed the CFHF diet was distinct from that of their WT littermates, while this difference was not present in the Control diet. 5xFAD mice on the CFHF diet had increased levels of Firmicutes and reduced levels of *Akkermansia municiphila* compared to WT mice on the same diet. The MCT diet had less of an effect on both the WT and 5xFAD microbiomes, but 5xFAD mice were more responsive. Indeed, 5xFAD mice on the MCT diet showed reduced alpha diversity due to reduced levels of firmicutes (*Lachnospiraceae, Oscillospiraceae, Lactobacillaceae, Ruminococcaceae*). These responses suggest that genetic mutations in AD alter the effects of ketogenic interventions on peripheral metabolism and microbiome.

### Limits of the study

A first limitation of this study is the sample size 3 or 4 per group, which if increased may reduce variability within the groups. Second, the lack of information regarding the levels of microbial metabolites limits the complete interpretation of microbiome alterations, as changes in the relative abundance of bacteria do not necessarily involve changes in metabolites. Similarly, 16S rRNA sequencing limits the interpretation of functional changes. A metatranscriptomic or metagenomic analysis would give more indication about the presence of the species, but also about their activity levels through the transcripts.

## CONCLUSION

This study reinforces the idea of microbial alterations existing in the early stages of AD. This suggests that genetic predispositions to AD induce a microbial shift toward lactate-producing bacteria and impaired metabolic processing of macronutrients. These alterations can be prevented by ketogenic interventions, which increase the levels of SCFA-producing bacteria and decrease those of the main lactate-producing bacteria. However, ketogenic strategies drive different bacterial genera and different mechanisms, which require further investigation. This study proposes that the microbiome might contribute to AD pathophysiology and represent a therapeutic target of ketogenic interventions.

## MATERIAL G METHODS

### Mice and treatment

#### Ethics

All experiments were carried out in accordance with the guidelines of the Canadian Council of Animal Care and were approved by the Institutional Animal Care Committee of the University of Sherbrooke.

#### 5xFAD mice

The generation and characterization of the model were previously described ^37,81^. The 5xFAD (Jackson laboratory MMRRC stock#: 34840-JAX) transgenic mouse model carries five familial Alzheimer’s disease (FAD) mutations, including **Swedish (K670N, M671L), Florida (I716V), and London (V717I)** mutations in the APP transgene, as well as **M146L and L286V FAD** mutations in the PSEN1 transgene, both under mouse Thy1 promoter. These transgenic mice were bred onto the B6SJLF1/J background (Jackson laboratory stock#: 100012). For our experiments in this study, we utilized both heterozygous mutant mice and non-carrier mice as appropriate controls.

#### 3xTg-AD mice and BC;12S strain wild type

The generation and characterization of the model were previously described^38,82^. 3xTg-AD mice (Jackson laboratory MMRC stock#: 34830-JAX) possess three human dementia-causing mutations that lead to AD’s pathological traits: **APP_Swe_ and tau_P301L_** encoded in two independent transgene constructs under control of the central nervous system (CNS) mouse regulatory element Thy1.2, co-microinjected into single-cell embryos harvested from homozygous mutant **PS1_M146V_** knock in (PS1-KI) mice. Wild type mice are B6;129 (WT, the 3xTg-AD background strain, Jackson laboratory stock#: 101045) obtained from a cross between C57BL/6J females (B6) and 129S1/SvImJ males (129S).

### Housing and Dietary interventions

Mice received either a Control diet (C, 70% carbohydrate, 20% long-chain triglycerides, 10% protein, 4.1 kcal/g, Research Diets product #D17121209I), a Control diet enriched with medium-chain triglycerides (MCT, 70% carbohydrate, 10% long-chain triglycerides, 10% medium-chain triglycerides (Caprylic acid to Capric acid ratio of 3:2), 10% protein, 4.1 kcal/g, Research Diets product #D17121210I) or a carbohydrate-free ketogenic-like diet (CFHF, 90% long-chain triglycerides, 10% protein, 6.7 kcal/g, Research Diets product# : D10070801). See table - detailed composition.

All animals were group-housed (3-4 per cage) in reverse cycle room (12h Dark-light cycle 10AM – OFF, 10PM – ON) with food/water ad libitum. 5xFAD mice were on diets for three months, spanning ages 2 to 5 months. 3xTg-AD mice were on diets for five months, spanning ages 3 to 8 months to analyze protective changes at pre-symptomatic level.

### Metabolic features

#### Body weight monitoring

The food pellets were changed twice a week according to the food company recommendations. The mice were weighed 28 days before the beginning of the experiment and twice from day 0 to day 28. From this date until day 140, mice were weighted once or twice a month.

### Beta-hydroxybutyrate monitoring

At different time points (Day0, Day14, Day85, Day140) circulating beta-hydroxybutyrate (BHB) levels were measured the freestyle precision neo (Abbott, product# : ART30531) with the blood BHB test strips (Abbott, REF# : 70748-75). The mice were pricked in the tail vein with a needle to have the required blood drops for the strips.

### Sample collection

Fecal matters were collected from each mouse at different timepoints of dietary interventions: Day 28, Day 85 and Day 140 (only 3xTg-AD mice) on diet. For 3xTg-AD mice, this corresponded to 4 months old, 6 months old and 8 months old. For 5xFAD mice, it was at 3 months old and 5 months old. The sample sizes were as follows:

#### Day 28 on diet

Control diet: WT n=4, 3xTg-AD n=4 / WT n=4, 5xFAD n=4

MCT diet: WT n=4, 3xTg-AD n=4 / WT n=4, 5xFAD n=4

CFHF diet: WT n=4, 3xTg-AD n=4 / WT n=4, 5xFAD n=4

#### Day 85 on diet

Control diet: WT n=4, 3xTg-AD n=4 / WT n=4, 5xFAD n=4

MCT diet: WT n=4, 3xTg-AD n=4 / WT n=4, 5xFAD n=4

CFHF diet: WT n=4, 3xTg-AD n=4 / WT n=4, 5xFAD n=4

#### Day 140 on diet

Control diet: WT n=1 (due to technical issues), 3xTg-AD n=4

MCT diet: WT n=3, 3xTg-AD n=3

CFHF diet: WT n=3, 3xTg-AD n=3

### DNA library preparation and 16S rRNA sequencing

16S amplicon libraries were prepared according to our previous works^83,84^. Extractions of microbial DNA from the stool samples were performed using the QIAamp DNA stool mini kit (Qiagen) with some modifications^85–87^. Briefly, stool sample is resuspended in 500 µl of ASL Buffer (Qiagen) with 250 µl of 0.1 mm glass beads. Samples were homogenized for 3 min at 6500 rpm (Precellys 24), then incubated at 95°C for 15min. Another round of homogenization was performed for 3 min at 6500 rpm and then centrifuged for 10 min at 13 000 rpm at room temperature. The supernatants were transferred to 1ml of 100 % EtOH and 10µl 3M sodium acetate (pH=5.2), then precipitated at -80°C for 15 min. Samples were centrifuged at 4°C for 20 min at 13 000 rpm and pellets were resuspended in 200 µl Tris 10 mM pH=7.5 and EDTA 1 mM. 2 µl for DNAse-Free RNase A (10 mg/mL) was added and incubated at 37°C for 30 min followed by the addition of 15 µl of proteinase K and 200µl of AL Buffer (Qiagen) incubated at 70°C for 30 min. 200 µl of 100% EtOH was added and transferred to a QIAamp column. The next step was done according to the manufacturer’s instructions and the DNA samples were stored at -80°C. For the amplification of the V4 region of the bacterial 16SrRNA gene, the 515F (5′-GTGCCAGCMGCCGCGGTAA-3′) and 806R (5′-GGACTACHVGGGTWTCTAAT-3′) paired primers were used with 5 ng of microbial genomic DNA^88^. Lastly, the pooled and indexed libraries (60 ng) were sequenced in paired-end modus on an Illumina MiSeq^89^ at the RNomics Platform of the University of Sherbrooke (https://rnomics.med.usherbrooke.ca/).

### Bioinformatics analysis

Quality control, filtering and trimming (truncate R1 and R2 reads after 240 and 160 bp, respectively; after truncation, sequences with more than 0 Ns and with higher than 2 "expected errors" were discarded; truncate reads at the first instance of a quality score less than or equal to 2) of fastq files, evaluation of error rate of amplicons, merging of paired reads, amplicon sequence variant table (ASV) table construction, chimera removal, taxonomy assigning (through SILVA database “silva nr99 v138.1 wSpecies train set.fa.gz”) were all performed with R package DADA2 v1.24.0^90^.

The overall structure of datasets and individual phenotypic characteristics were explored though multidimensional analysis plots such as heatmap (from the 200 most variable ASVs) and multidimensional scaling (MDS) plots utilizing R package metagenomeSeq v1.38.0^91,92^. The first 3 most important MDS axes where 3D visualized through R package compositions v2.0-8. Within-habitat diversity species richness (Alpha diversity) analysis was performed through R package phyloseq v1.30.0^93^ utilizing Shannon, Simpson and Chao1 indices. Mean comparisons were tested through R package ggpubr v0.6.0 and included non-parametric Wilcoxon test for two-group comparison and non-parametric Kruskal-Wallis test for multiple groups comparisons. Variance in the structure of the population (Beta diversity) with Bray-Curtis metric was also performed with phyloseq v1.30.0. Here to test if the groups were significantly different from each other we performed a Permanova test using the adonis function from the vegan v2.6-4 R package. To track the longitudinal microbiome changes, more specifically to check how beta diversity changed across diet time, we created volatility plots through QIIME2^94^ using Unweighted Unifrac and Jaccard metrics.

Next, we performed a differential expression analysis to statistically compare previously obtained ASVs among the different pre-determined groups (WT C vs 3xTg C, WT C vs 3xTg M, WT C vs 3xTg CFHF). First, due to varying depths of coverage across mice, we needed to normalize data. To reduce biases resulting from preferentially sampled taxa we normalized by using CSS (Cumulative-Sum Scaling)^95^ implemented in metagenomeSeq^91^. We sampled same mice between different conditions, so their duplicated effect cannot be omitted from the successive statistical analysis, however differences between mice were not of direct interest; to account for all of this we decided to treat mice IDs as random effect in a mixed effects model where our fixed effects were just reduced to our group of interest^96^. Thus, mice IDs were considered as a blocking variable. Their apport was calculated though the function duplicateCorrelation of R package limma v3.50.1^97^ and included in the subsequent fitted model. All interesting contrasts (WT C vs 3xTg C, WT C vs 3xTg M, WT C vs 3xTg CFHF) were then statistically evaluated and visualized (through volcano plots) with limma where differential expressed ASVs were filtered at a p-adjusted threshold value at 0.05. Heatmaps were also used to visualize the diet effect on microbial alterations of AD mice. They were generated with R package Pretty Heatmaps (Raivo Kolde (2019). pheatmap: Pretty Heatmaps. R package version 1.0.12. https://CRAN.R-project.org/package=pheatmap).

Venn diagrams were created to better visualize common differentially expressed ASVs among conditions of interest. The same was done for 5xFAD mice following the interesting contrasts (WT C vs 5xFAD C, WT C vs 5xFAD M, WT C vs 5xFAD CFHF, WT C vs WT M, 5xFAD C vs 5xFAD M, WT C vs WT CFHF, 5xFAD C vs 5xFAD CFHF).

To predict functional abundances based on marker gene sequences we used PICRUSt2^98^. We then performed differential abundance (DA) analysis, using ALDEx2 Kruskal-Wallace test, on its abundance data outputs, annotated under Metacyc and KEGG consortia, with R package ggpicrust2 v1.7.330^99^. We filter out those pathways that did not passed a p-value adjusted of 0.05. Using resulting differentially expressed pathways we plotted a heatmap (R package Pretty Heatmaps) and barplots with error bars (to limit the results to better visualize the plot, here we ordered by decrescent adjusted p-value and selected just the top 20). Metacyc pathway PCA was performed with R package factoextra v1.0.7.

### Statistical analysis

The correlation between varying ASVs and predicted functional changes was calculated in R using Pearson method. We filtered out the correlation that did not passed an adjusted p-value of 0.05 (Benjamini and Hochberg’s method) and visualized with ggplot heatmap. Statistical tests for metabolic monitoring and relative abundance levels at phylum and family levels were performed using GraphPad Prism v10.0(GraphPad Software). Mixed effect analysis and two-way ANOVA with Uncorrected Fisher’s LSD test were used for appropriate situations. The pair-wise comparison tests were performed only for the following contrasts : for strain effect within the same diet/ WT C vs 3xTg C, WT M vs 3xTg M, WT CFHF vs 3xTg CFHF, for diet effect within the same strain / WT C vs WT M, WT C vs WT CFHF, 3xTg C vs 3xTg M, 3xTg C vs 3xTg CFHF. Data are presented as means ± standard errors of the mean. Statistical tests performed are detailed in the legend for each figure. P-values less than 0.05 were considered statistically significant and p-values between 0.05 and 0.1 were considered nearly significant.

### Nomenclature

A specific bacterium is named based on its genus and species (ex: *Bifidobacterium pseudolongum*). When the specific genus or the species is not known, the bacterium is named with the most precis taxonomical group plus “sp.” in the text or “NA” on the graphs (ex: *Muribaculaceae sp.*, *Muribaculaceae_NA_NA)*.

## Data availability

The data underlying this article are available in the article and in its online supplementary material. Sequencing data are available on Sequence Read Archive (SRA) : PRJNA1160852

## AUTHOR CONTRIBUTIONS

P.E.H.M developed the concept, carried out the experiments, collected samples, analyzed data, and wrote the manuscript. M.A performed bioinformatic analysis. E.N.K.L and K.P. extracted DNA and supervised 16S rRNA sequencing. M.M. and A.A. collected samples and performed metabolic monitoring. P.B. analyzed data. E.M. supervised 16S rRNA sequencing, analyzed data and revised the manuscript. K.J.L.F developed the concept, analyzed data, and revised the manuscript. All authors read and approved the final manuscript.

## ACKNOWLEDGEMENT

We thank the platform of transcriptomic analysis (“RNomique”) of the Faculté de Médecine et des Sciences de la Santé at the Université de Sherbrooke for the 16S rRNA sequencing. This work was funded by operating grants to K.J.L.F. from the Canadian Institutes of Health Research (CIHR), the Natural Sciences and Engineering Research Council (NSERC), and a Tier 1 Canada Research Chair. E.M. holds grant from the CIHR [MOP389354]. P.E.H.M. was supported by the Bourse d’exemption from the Department of Neurosciences at the Université de Montréal.

## Supplementary information

**Supplementary figure 1 –** related to Figure 1

**Supplementary figure 2 & 3 –** related to Figure 2

**Supplementary figure 4 –** related to Figure 3

**Supplementary figure 5 –** related to Figure 4

**Supplementary figure 6 –** related to Figure 3 & 4

**Supplementary figure 7 –** related to Figure 4

**Supplementary figure 8 –** related to Figure 5

**Supplementary table 1 –** Relative abundance of each varying ASV in 3xTg-AD mice after 1, 3 or 5 months of ketogenic interventions in 3xTg-AD mice – related to Figure 3

**Supplementary Table 2 –** Relative abundance of each varying ASV in 5xFAD mice after 1 month and 3 months of ketogenic interventions – related to Supp. Figure 4

**Supplementary table 3 –** Correlation matrix between varying ASVs and predicted altered Metacyc pathways after 1 month of ketogenic interventions in 3xTg-AD mice – related to Figure 5

**Supplementary Figure 1 - Monitoring of metabolic features and overview of faecal microbiome features in 5xFAD mice treated with ketogenic interventions**

**a)** Body weight monitoring in mice fed diets(n=6-7/strain/diet). The mixed-effect stats model is based on following fixed effects: Time on diet (p-value<0.0001, F(3.934, 145.6) = 104.5), Strain X Diet (p-value=n.s.), Interaction (Time x Strain X Diet, p-value=0.0001, F (60, 444) = 1.916).

The arrows indicate the start of the diet (black) and the time when fecal matter was harvested (brown).

**b)** Blood levels of beta-hydroxybutyrate(BHB) at different timepoints. The 2-way ANOVA is based on two variables: Strain and Diet.

Day 0(D0): Strain (p-value = n.s.), Diet (p-value=n.s.), Interaction(p-value=n.s.);

Day14(D14): Strain(p-value =n.s.), Diet (p-value<0.0001, F(2, 37) = 105.5), Interaction(p-value=n.s., F(2, 37) = 0.2839);

Day85(D85): Strain (p-value =n.s.),Diet(p-value<0.0001, F (2, 37) = 29.27),Interaction(p-value=0.041, F(2, 37) = 3.473)

**c)** Multidimensional scaling (MDS) analysis of microbiome of experimental groups(n=4/strain/diet/timepoint) with percentage of variance explained, legend indicate strain, diet and time

**d)** Unbiased heatmap of all varying bacterial species abundance in fecal microbiome with hierarchical clustering of rows (218 ASVs) and columns (48 samples, n=4/strain/diet/timepoint); legend indicates strain, diet, and level of abundance.

Significant multiple comparisons (uncorr. Fisher’s LSD) on graph: : *p-value<0.05; **p-value<0.01; ***p-value<0.001; ****p-value<0.0001

**Supplementary Figure 2 -Longitudinal assessment of Alpha and Beta diversity of the microbiome in 5xFAD mice**

a-c)Indices of Alpha diversity at different time points of the diet including a)Shannon, b)Simpson, and c)Chao1 indices(multivariate statistical analysis with Kruskal Wallis and pair-wise comparison post-hoc test: Wilcoxon, resuts on graph).

d-f) Indices of Beta diversity including d)Bray-Curtis showing PCoA1 and PCoA2 (Multivariate statistical analysis with Adonis2 considering the following factors : group(strainXdiet) and residual, results for group factor on graph),

e) Volatility plot of the average changes in PCoA1 of the Unweighted Unifrac distance matrix,and f) Volatility plot of the average changes in PCoA1 of the Jaccard distance matrix.

**Supplementary Figure 3 – Evaluation of time effect on the Alpha diversity within the microbiome of 3xTg-AD and 5xFAD mice treated with ketogenic interventions**

**a-c)** Analysis of the significance of time factor for Alpha-diversity indices including a)Shannon, b)Simpson, c)Chao1 indices in 3xTg-AD mice (Multivariate statistical analysis with Kruskal Wallis and pair-wise comparison post-hoc test : Wilcoxon, resuts on graph)

**d-f)** Analysis of the significance of time factor for Alpha-diversity indices including **d)**Shannon, **e)**Simpson, **f)**Chao1 indices in 5xFAD mice (Multivariate statistical analysis with Kruskal Wallis and pair-wise comparison post-hoc test : Wilcoxon, resuts on graph)

**Supplementary Figure 4 – Diet-induced alterations of relative abundance levels after 1 month and 3 months intervention in 5xFAD mice**

**a-d)** Differential expression analysis after 1 month of ketogenic interventions, obtained with Limma (adjusted p-value <0.05) to **a)** visualize significant changes in relative abundance levels of ASVs between WT and 5xFAD mice on Control diet (WT C vs 5xFAD C) with Volcano plot;

**b)** Visualize ASVs restored by the MCT or CFHF diet with a heatmap. The ASVs that are significantly different in WT C vs 5xFAD C and not different in WT C vs 5xFAD M or WT C vs 5xFAD CFHF are considered “restored”; visualize with hierarchical heatmaps the new bacterial changes induced by **c)** MCT and **d)** CFHF diet in 5xFAD mice. The color code in the extreme left column indicates whether the ASV is altered in WT mice (cyan), 5xFAD mice (pink), or in both genotypes (purple).

**e-f)** Differential expression analysis after 3 months of ketogenic interventions, obtained with Limma (adjusted p-value <0.05) showing hierarchical heatmaps of the new bacterial changes induced by **e)** MCT and **f)** CFHF diet in 5xFAD mice. The color code in the extreme left column is the same as **Supp.Fig. 4c-d**

**Supplementary Figure 5 – Changes in the relative abundance levels of bacteria after 1 month of ketogenic interventions at Phylum and Family levels in 5xFAD mice**

**a)** Relative abundance levels of identified phyla alterations (Stats on graph : For each phylum - 2way-ANOVA based on Strain and Diet factors and Uncorrected Fisher’s LSD for pair-wise comparison tests),

**b)** Ratio of *Firmicutes:Bacteroidetes* (F/B ratio, Stats on graph : 2way-ANOVA based on Strain and Diet factors and Uncorrected Fisher’s LSD for pair-wise comparison tests),

**c)** Relative abundance of the *Firmicutes* families showing Strain or Diet effect (Stats on graph: For each family: 2way-ANOVA based on Strain and Diet factors and Uncorrected Fisher’s LSD for pair-wise comparison tests).

**d)** Relative abundance of the *Proteobacteria* families showing Strain or Diet effect (Stats on graph: For each family: 2way-ANOVA based on Strain and Diet factors and Uncorrected Fisher’s LSD for pair-wise comparison tests).

Stats : *p-value<0.05; **p-value<0.01; ***p-value<0.001; ****p-value<0.0001

**Supplementary Figure 6 – Changes in the relative abundance levels of firmicutes families and top 20 predicted Metacyc pathways alterations after 1 month of ketogenic interventions in 3xTg-AD mice**

**a)** Venn diagram comparing the number of ASVs altered in 3xTg AD mice on Control diet at 4 months old (WTC vs 3xTgC 4mo) to the number of ASVs altered in 3xTg AD mice on Control diet at 6 months old (WTC vs 3xTgC 6mo). 83% of ASVs altered at 6mo are already present at 4 months old. Analysis performed at 4 months of age or after 1 month of ketogenic interventions is still relevant for 6 months old mice.

**b)** Relative abundance of the *Firmicutes* families showing Strain or Diet effect (Stats on graph : For each family : 2way-ANOVA based on Strain and Diet factors and Uncorrected Fisher’s LSD for pair-wise comparison tests).

**c)** Top 20 predicted Metacyc pathways with their relative abundance and adjusted p-value (ALDEx2 Kruskal-Wallace)

**Supplementary Figure 7 – Predicted KEGG pathways alterations with their relative abundance after 1 month of ketogenic interventions**

**Supplementary Figure 8 – Correlation between varying ASVs and predicted Metacyc pathways decreased or not affected in 3xTg-AD mice on Control diet but increased by ketogenic interventions**

This heatmap shows the remaining pathways after filtering out ketone or SCFA-related pathways.

## References

1. Mosconi, L., Brys, M., Glodzik-Sobanska, L., De Santi, S., Rusinek, H., and de Leon, M.J. (2007). Early detection of Alzheimer’s disease using neuroimaging. Exp Gerontol 42, 129–138. 10.1016/j.exger.2006.05.016.

2. Association, A.s. (2021). 2021 Alzheimer’s disease facts and figures. Alzheimer’s & Dementia 17, 327–406. 10.1002/alz.12328.

3. Arnold, S.E., Arvanitakis, Z., Macauley-Rambach, S.L., Koenig, A.M., Wang, H.-Y., Ahima, R.S., Craft, S., Gandy, S., Buettner, C., Stoeckel, L.E., et al. (2018). Brain insulin resistance in type 2 diabetes and Alzheimer disease: concepts and conundrums. Nature Reviews Neurology 14, 168–181. 10.1038/nrneurol.2017.185.

4. Livingston, G., Huntley, J., Sommerlad, A., Ames, D., Ballard, C., Banerjee, S., Brayne, C., Burns, A., Cohen-Mansfield, J., Cooper, C., et al. (2020). Dementia prevention, intervention, and care: 2020 report of the *Lancet* Commission. The Lancet 396, 413–446. 10.1016/S0140-6736(20)30367-6.

5. Scheltens, P., De Strooper, B., Kivipelto, M., Holstege, H., Chételat, G., Teunissen, C.E., Cummings, J., and van der Flier, W.M. (2021). Alzheimer’s disease. The Lancet 397, 1577–1590. 10.1016/S0140-6736(20)32205-4.

6. Trushina, E. (2019). Alzheimer’s disease mechanisms in peripheral cells: Promises and challenges. Alzheimers Dement (N Y) 5, 652–660. 10.1016/j.trci.2019.06.008.

7. Fortier, M., Castellano, C.-A., St-Pierre, V., Myette-Côté, É., Langlois, F., Roy, M., Morin, M.-C., Bocti, C., Fulop, T., Godin, J.-P., et al. (2020). A ketogenic drink improves cognition in mild cognitive impairment: Results of a 6-month RCT. Alzheimer’s & Dementia n/a. 10.1002/alz.12206.

8. Phillips, M.C.L., Deprez, L.M., Mortimer, G.M.N., Murtagh, D.K.J., McCoy, S., Mylchreest, R., Gilbertson, L.J., Clark, K.M., Simpson, P.V., McManus, E.J., et al. (2021). Randomized crossover trial of a modified ketogenic diet in Alzheimer’s disease. Alzheimer’s Research & Therapy 13, 51. 10.1186/s13195-021-00783-x.

9. Newman, J.C., and Verdin, E. (2017). β-Hydroxybutyrate: A Signaling Metabolite. Annu Rev Nutr 37, 51–76. 10.1146/annurev-nutr-071816-064916.

10. Augustin, K., Khabbush, A., Williams, S., Eaton, S., Orford, M., Cross, J.H., Heales, S.J.R., Walker, M.C., and Williams, R.S.B. (2018). Mechanisms of action for the medium-chain triglyceride ketogenic diet in neurological and metabolic disorders. Lancet Neurol 17, 84–93. 10.1016/s1474-4422(17)30408-8.

11. Vandenberghe, C., St-Pierre, V., Fortier, M., Castellano, C.-A., Cuenoud, B., and Cunnane, S.C. (2020). Medium Chain Triglycerides Modulate the Ketogenic Effect of a Metabolic Switch. Frontiers in Nutrition 7. 10.3389/fnut.2020.00003.

12. Lin, T.Y., Liu, H.W., and Hung, T.M. (2021). The Ketogenic Effect of Medium-Chain Triacylglycerides. Front Nutr 8, 747284. 10.3389/fnut.2021.747284.

13. Grammatikopoulou, M.G., Goulis, D.G., Gkiouras, K., Theodoridis, X., Gkouskou, K.K., Evangeliou, A., Dardiotis, E., and Bogdanos, D.P. (2020). To Keto or Not to Keto? A Systematic Review of Randomized Controlled Trials Assessing the Effects of Ketogenic Therapy on Alzheimer Disease. Advances in Nutrition 11, 1583–1602. 10.1093/advances/nmaa073.

14. Cunnane, S.C., Trushina, E., Morland, C., Prigione, A., Casadesus, G., Andrews, Z.B., Beal, M.F., Bergersen, L.H., Brinton, R.D., de la Monte, S., et al. (2020). Brain energy rescue: an emerging therapeutic concept for neurodegenerative disorders of ageing. Nat Rev Drug Discov 19, 609–633. 10.1038/s41573-020-0072-x.

15. Castro, C.B., Dias, C.B., Hillebrandt, H., Sohrabi, H.R., Chatterjee, P., Shah, T.M., Fuller, S.J., Garg, M.L., and Martins, R.N. (2023). Medium-chain fatty acids for the prevention or treatment of Alzheimer’s disease: a systematic review and meta-analysis. Nutrition Reviews 81, 1144–1162. 10.1093/nutrit/nuac104.

16. Newman, J.C., and Verdin, E. (2014). Ketone bodies as signaling metabolites. Trends Endocrinol Metab 25, 42–52. 10.1016/j.tem.2013.09.002.

17. Mazandarani, M., Lashkarbolouk, N., Ejtahed, H.-S., and Qorbani, M. (2023). Does the ketogenic diet improve neurological disorders by influencing gut microbiota? A systematic review. Nutrition Journal 22, 61. 10.1186/s12937-023-00893-2.

18. Khedr, E.M., Omeran, N., Karam-Allah Ramadan, H., Ahmed, G.K., and Abdelwarith, A.M. (2022). Alteration of Gut Microbiota in Alzheimer’s Disease and Their Relation to the Cognitive Impairment. J Alzheimers Dis 88, 1103–1114. 10.3233/jad-220176.

19. Vogt, N.M., Kerby, R.L., Dill-McFarland, K.A., Harding, S.J., Merluzzi, A.P., Johnson, S.C., Carlsson, C.M., Asthana, S., Zetterberg, H., Blennow, K., et al. (2017). Gut microbiome alterations in Alzheimer’s disease. Scientific Reports 7, 13537. 10.1038/s41598-017-13601-y.

20. Liu, P., Wu, L., Peng, G., Han, Y., Tang, R., Ge, J., Zhang, L., Jia, L., Yue, S., Zhou, K., et al. (2019). Altered microbiomes distinguish Alzheimer’s disease from amnestic mild cognitive impairment and health in a Chinese cohort. Brain, Behavior, and Immunity 80, 633–643. 10.1016/j.bbi.2019.05.008.

21. Lopizzo, N., Provasi, S., Marizzoni, M., Borruso, L., Andryszak, P., Frisoni, G.B., and Cattaneo, A. (2019). Identification of gut microbiota signature in Alzheimer’s disease: Possible role in influencing peripheral inflammation. European Neuropsychopharmacology 29, S167–S168. 10.1016/j.euroneuro.2018.11.289.

22. Cattaneo, A., Cattane, N., Galluzzi, S., Provasi, S., Lopizzo, N., Festari, C., Ferrari, C., Guerra, U.P., Paghera, B., Muscio, C., et al. (2017). Association of brain amyloidosis with pro-inflammatory gut bacterial taxa and peripheral inflammation markers in cognitively impaired elderly. Neurobiology of Aging 49, 60–68. 10.1016/j.neurobiolaging.2016.08.019.

23. Borsom, E.M., Conn, K., Keefe, C.R., Herman, C., Orsini, G.M., Hirsch, A.H., Avila, M.P., Testo, G., Jaramillo, S.A., Bolyen, E., et al. (2023). Predicting Neurodegenerative Disease Using Prepathology Gut Microbiota Composition: a Longitudinal Study in Mice Modeling Alzheimer’s Disease Pathologies. Microbiology Spectrum 11, e03458–03422. doi:10.1128/spectrum.03458-22.

24. Laske, C., Müller, S., Preische, O., Ruschil, V., Munk, M.H.J., Honold, I., Peter, S., Schoppmeier, U., and Willmann, M. (2022). Signature of Alzheimer’s Disease in Intestinal Microbiome: Results From the AlzBiom Study. Frontiers in Neuroscience 16. 10.3389/fnins.2022.792996.

25. Bairamian, D., Sha, S., Rolhion, N., Sokol, H., Dorothée, G., Lemere, C.A., and Krantic, S. (2022). Microbiota in neuroinflammation and synaptic dysfunction: a focus on Alzheimer’s disease. Molecular Neurodegeneration 17, 19. 10.1186/s13024-022-00522-2.

26. Shabbir, U., Arshad, M.S., Sameen, A., and Oh, D.-H. (2021). Crosstalk between Gut and Brain in Alzheimer’s Disease: The Role of Gut Microbiota Modulation Strategies. Nutrients 13, 690.

27. Zhang, T., Gao, G., Kwok, L.-Y., and Sun, Z. (2023). Gut microbiome-targeted therapies for Alzheimer’s disease. Gut Microbes 15, 2271613. 10.1080/19490976.2023.2271613.

28. Du, Y., He, C., An, Y., Huang, Y., Zhang, H., Fu, W., Wang, M., Shan, Z., Xie, J., Yang, Y., and Zhao, B. (2024). The Role of Short Chain Fatty Acids in Inflammation and Body Health. International Journal of Molecular Sciences 25, 7379.

29. Kimura, I., Inoue, D., Maeda, T., Hara, T., Ichimura, A., Miyauchi, S., Kobayashi, M., Hirasawa, A., and Tsujimoto, G. (2011). Short-chain fatty acids and ketones directly regulate sympathetic nervous system via G protein-coupled receptor 41 (GPR41). Proceedings of the National Academy of Sciences 108, 8030–8035. 10.1073/pnas.1016088108.

30. Dalile, B., Van Oudenhove, L., Vervliet, B., and Verbeke, K. (2019). The role of short-chain fatty acids in microbiota–gut–brain communication. Nature Reviews Gastroenterology & Hepatology 16, 461–478. 10.1038/s41575-019-0157-3.

31. Fusco, W., Lorenzo, M.B., Cintoni, M., Porcari, S., Rinninella, E., Kaitsas, F., Lener, E., Mele, M.C., Gasbarrini, A., Collado, M.C., et al. (2023). Short-Chain Fatty-Acid-Producing Bacteria: Key Components of the Human Gut Microbiota. Nutrients 15. 10.3390/nu15092211.

32. Ling, Z., Zhu, M., Yan, X., Cheng, Y., Shao, L., Liu, X., Jiang, R., and Wu, S. (2021). Structural and Functional Dysbiosis of Fecal Microbiota in Chinese Patients With Alzheimer’s Disease. Frontiers in Cell and Developmental Biology 8. 10.3389/fcell.2020.634069.

33. Sholl, J., Mailing, L.J., and Wood, T.R. (2021). Reframing Nutritional Microbiota Studies To Reflect an Inherent Metabolic Flexibility of the Human Gut: a Narrative Review Focusing on High-Fat Diets. mBio 12, 10.1128/mbio.00579-00521. doi:10.1128/mbio.00579-21.

34. Gong, X., Cai, Q., Liu, X., An, D., Zhou, D., Luo, R., Peng, R., and Hong, Z. (2021). Gut flora and metabolism are altered in epilepsy and partially restored after ketogenic diets. Microb Pathog 155, 104899. 10.1016/j.micpath.2021.104899.

35. Rew, L., Harris, M.D., and Goldie, J. (2022). The ketogenic diet: its impact on human gut microbiota and potential consequent health outcomes: a systematic literature review. Gastroenterol Hepatol Bed Bench 15, 326–342. 10.22037/ghfbb.v15i4.2600.

36. Nagpal, R., Neth, B.J., Wang, S., Craft, S., and Yadav, H. (2019). Modified Mediterranean-ketogenic diet modulates gut microbiome and short-chain fatty acids in association with Alzheimer’s disease markers in subjects with mild cognitive impairment. EBioMedicine 47, 529–542. 10.1016/j.ebiom.2019.08.032.

37. Oakley, H., Cole, S.L., Logan, S., Maus, E., Shao, P., Craft, J., Guillozet-Bongaarts, A., Ohno, M., Disterhoft, J., Eldik, L.V., et al. (2006). Intraneuronal β-Amyloid Aggregates, Neurodegeneration, and Neuron Loss in Transgenic Mice with Five Familial Alzheimer’s Disease Mutations: Potential Factors in Amyloid Plaque Formation. The Journal of Neuroscience 26, 10129–10140. 10.1523/jneurosci.1202-06.2006.

38. Oddo, S., Caccamo, A., Shepherd, J.D., Murphy, M.P., Golde, T.E., Kayed, R., Metherate, R., Mattson, M.P., Akbari, Y., and LaFerla, F.M. (2003). Triple-transgenic model of Alzheimer’s disease with plaques and tangles: intracellular Abeta and synaptic dysfunction. Neuron 39, 409–421. 10.1016/s0896-6273(03)00434-3.

39. Peeters, J., Thas, O., Shkedy, Z., Kodalci, L., Musisi, C., Owokotomo, O.E., Dyczko, A., Hamad, I., Vangronsveld, J., Kleinewietfeld, M., et al. (2021). Exploring the Microbiome Analysis and Visualization Landscape. Frontiers in Bioinformatics 1. 10.3389/fbinf.2021.774631.

40. Wang, Y., Xiao, J., Wei, S., Su, Y., Yang, X., Su, S., Lan, L., Chen, X., Huang, T., and Shan, Q. (2024). Protective effect of zinc gluconate on intestinal mucosal barrier injury in antibiotics and LPS-induced mice. Frontiers in Microbiology 15. 10.3389/fmicb.2024.1407091.

41. Bliek Tijs, F.v.d.K., Marc Galland. RNA-seq lesson. https://github.com/ScienceParkStudyGroup/rnaseq-lesson.

42. Chen, J., Bittinger, K., Charlson, E.S., Hoffmann, C., Lewis, J., Wu, G.D., Collman, R.G., Bushman, F.D., and Li, H. (2012). Associating microbiome composition with environmental covariates using generalized UniFrac distances. Bioinformatics 28, 2106–2113. 10.1093/bioinformatics/bts342.

43. Smith, E., and Morowitz, H.J. (2004). Universality in intermediary metabolism. Proceedings of the National Academy of Sciences 101, 13168–13173. doi:10.1073/pnas.0404922101.

44. Caspi, R., Billington, R., Keseler, I.M., Kothari, A., Krummenacker, M., Midford, P.E., Ong, W.K., Paley, S., Subhraveti, P., and Karp, P.D. (2020). The MetaCyc database of metabolic pathways and enzymes - a 2019 update. Nucleic Acids Res 48, D445–d453. 10.1093/nar/gkz862.

45. O’Callaghan, A., and van Sinderen, D. (2016). Bifidobacteria and Their Role as Members of the Human Gut Microbiota. Frontiers in Microbiology 7. 10.3389/fmicb.2016.00925.

46. Ross, F.C., Patangia, D., Grimaud, G., Lavelle, A., Dempsey, E.M., R, P.R., and Stanton, C. (2024). The interplay between diet and the gut microbiome: implications for health and disease. Nature Reviews Microbiology, 1-16. 10.1038/s41579-024-01068-4.

47. Ayten, Ş., and Bilici, S. (2024). Modulation of Gut Microbiota Through Dietary Intervention in Neuroinflammation and Alzheimer’s and Parkinson’s Diseases. Current Nutrition Reports 13, 82–96. 10.1007/s13668-024-00539-7.

48. Ang, Q.Y., Alexander, M., Newman, J.C., Tian, Y., Cai, J., Upadhyay, V., Turnbaugh, J.A., Verdin, E., Hall, K.D., Leibel, R.L., et al. (2020). Ketogenic Diets Alter the Gut Microbiome Resulting in Decreased Intestinal Th17 Cells. Cell 181, 1263–1275.e1216. 10.1016/j.cell.2020.04.027.

49. Dilimulati, D., Zhang, F., Shao, S., Lv, T., Lu, Q., Cao, M., Jin, Y., Jia, F., and Zhang, X. (2023). Ketogenic Diet Modulates Neuroinflammation via Metabolites from Lactobacillus reuteri After Repetitive Mild Traumatic Brain Injury in Adolescent Mice. Cellular and Molecular Neurobiology 43, 907–923. 10.1007/s10571-022-01226-3.

50. Klein, M.S., Newell, C., Bomhof, M.R., Reimer, R.A., Hittel, D.S., Rho, J.M., Vogel, H.J., and Shearer, J. (2016). Metabolomic Modeling To Monitor Host Responsiveness to Gut Microbiota Manipulation in the BTBRT+tf/j Mouse. Journal of Proteome Research 15, 1143–1150. 10.1021/acs.jproteome.5b01025.

51. Vacca, M., Celano, G., Calabrese, F.M., Portincasa, P., Gobbetti, M., and De Angelis, M. (2020). The Controversial Role of Human Gut Lachnospiraceae. Microorganisms 8. 10.3390/microorganisms8040573.

52. You, H., Tan, Y., Yu, D., Qiu, S., Bai, Y., He, J., Cao, H., Che, Q., Guo, J., and Su, Z. (2022). The Therapeutic Effect of SCFA-Mediated Regulation of the Intestinal Environment on Obesity. Frontiers in Nutrition 9. 10.3389/fnut.2022.886902.

53. St-Pierre, V., Vandenberghe, C., Lowry, C.-M., Fortier, M., Castellano, C.-A., Wagner, R., and Cunnane, S.C. (2019). Plasma Ketone and Medium Chain Fatty Acid Response in Humans Consuming Different Medium Chain Triglycerides During a Metabolic Study Day. Frontiers in Nutrition 6. 10.3389/fnut.2019.00046.

54. Shcherbakova, K., Schwarz, A., Ivleva, I., Nikitina, V., Krytskaya, D., Apryatin, S., Karpenko, M., and Trofimov, A. (2023). Short- and long-term cognitive and metabolic effects of medium-chain triglyceride supplementation in rats. Heliyon 9, e13446. 10.1016/j.heliyon.2023.e13446.

55. Ota, M., Matsuo, J., Ishida, I., Takano, H., Yokoi, Y., Hori, H., Yoshida, S., Ashida, K., Nakamura, K., Takahashi, T., and Kunugi, H. (2019). Effects of a medium-chain triglyceride-based ketogenic formula on cognitive function in patients with mild-to-moderate Alzheimer’s disease. Neuroscience Letters 690, 232–236. 10.1016/j.neulet.2018.10.048.

56. Zhang, J., Yu, H., Wang, Q., Cai, H., Shen, F., Ruan, S., Wu, Y., Liu, T., Feng, F., and Zhao, M. (2023). Dietary additive octyl and decyl glycerate modulates metabolism and inflammation under different dietary patterns with the contribution of the gut microbiota. Food & Function 14, 525–540. 10.1039/D2FO03059D.

57. Zhang, W., Chen, S., Huang, X., Tong, H., Niu, H., and Lu, L. (2023). Neuroprotective effect of a medium-chain triglyceride ketogenic diet on MPTP-induced Parkinson’s disease mice: a combination of transcriptomics and metabolomics in the substantia nigra and fecal microbiome. Cell Death Discovery 9, 251. 10.1038/s41420-023-01549-0.

58. Zhou, J., Wang, J., Yao, M., He, J., Yang, Y., Li, X., Tan, Z., Shi, H., Zhu, X., and Tian, B. (2022). An acetate-independent pathway for isopropanol production via HMG-CoA in Escherichia coli. Journal of Biotechnology 359, 29–34. 10.1016/j.jbiotec.2022.09.011.

59. Jung, J.H., Kim, G., Byun, M.S., Lee, J.H., Yi, D., Park, H., Lee, D.Y., and for the, K.R.G. (2022). Gut microbiome alterations in preclinical Alzheimer’s disease. PLOS ONE 17, e0278276. 10.1371/journal.pone.0278276.

60. Li, B., He, Y., Ma, J., Huang, P., Du, J., Cao, L., Wang, Y., Xiao, Q., Tang, H., and Chen, S. (2019). Mild cognitive impairment has similar alterations as Alzheimer’s disease in gut microbiota. Alzheimer’s & Dementia 15, 1357–1366. 10.1016/j.jalz.2019.07.002.

61. Yu, B., Wan, G., Cheng, S., Wen, P., Yang, X., Li, J., Tian, H., Gao, Y., Zhong, Q., Liu, J., et al. (2024). Disruptions of Gut Microbiota are Associated with Cognitive Deficit of Preclinical Alzheimer’s Disease: A Cross-Sectional Study. Curr Alzheimer Res. 10.2174/0115672050303878240319054149.

62. Chen, G., Zhou, X., Zhu, Y., Shi, W., and Kong, L. (2023). Gut microbiome characteristics in subjective cognitive decline, mild cognitive impairment and Alzheimer’s disease: a systematic review and meta-analysis. European Journal of Neurology 30, 3568–3580. 10.1111/ene.15961.

63. Bonnechère, B., Amin, N., and van Duijn, C. (2022). What Are the Key Gut Microbiota Involved in Neurological Diseases? A Systematic Review. International Journal of Molecular Sciences 23, 13665.

64. Fabi, J.P. (2024). The connection between gut microbiota and its metabolites with neurodegenerative diseases in humans. Metabolic Brain Disease 39, 967–984. 10.1007/s11011-024-01369-w.

65. Zhuang, Z.-Q., Shen, L.-L., Li, W.-W., Fu, X., Zeng, F., Gui, L., Lü, Y., Cai, M., Zhu, C., Tan, Y.-L., et al. (2018). Gut Microbiota is Altered in Patients with Alzheimer’s Disease. Journal of Alzheimer’s Disease 63, 1337–1346. 10.3233/JAD-180176.

66. Haran John, P., Bhattarai Shakti, K., Foley Sage, E., Dutta, P., Ward Doyle, V., Bucci, V., and McCormick Beth, A. (2019). Alzheimer’s Disease Microbiome Is Associated with Dysregulation of the Anti-Inflammatory P-Glycoprotein Pathway. mBio 10, 10.1128/mbio.00632-00619. 10.1128/mbio.00632-19.

67. Chen, L., Xu, X., Wu, X., Cao, H., Li, X., Hou, Z., Wang, B., Liu, J., Ji, X., Zhang, P., and Li, H. (2022). A comparison of the composition and functions of the oral and gut microbiotas in Alzheimer’s patients. Front Cell Infect Microbiol 12, 942460. 10.3389/fcimb.2022.942460.

68. Jeong, S., Huang, L.-K., Tsai, M.-J., Liao, Y.-T., Lin, Y.-S., Hu, C.-J., and Hsu, Y.-H. (2022). Cognitive Function Associated with Gut Microbial Abundance in Sucrose and S-Adenosyl-L-Methionine (SAMe) Metabolic Pathways. Journal of Alzheimer’s Disease 87, 1115–1130. 10.3233/JAD-215090.

69. Wang, Y., Ma, M., Dai, W., Shang, Q., and Yu, G. (2024). Bacteroides salyersiae is a potent chondroitin sulfate-degrading species in the human gut microbiota. Microbiome 12, 41. 10.1186/s40168-024-01768-2.

70. He, K., Liu, M., Wang, Q., Chen, S., and Guo, X. (2023). Combined analysis of 16S rDNA sequencing and metabolomics to find biomarkers of drug-induced liver injury. Scientific Reports 13, 15138. 10.1038/s41598-023-42312-w.

71. Luo, Y.-X., Yang, L.-L., and Yao, X.-Q. (2024). Gut microbiota-host lipid crosstalk in Alzheimer’s disease: implications for disease progression and therapeutics. Molecular Neurodegeneration 19, 35. 10.1186/s13024-024-00720-0.

72. Lin, J.-z., Duan, M.-r., Lin, N., and Zhao, W.-j. (2021). The emerging role of the chondroitin sulfate proteoglycan family in neurodegenerative diseases. Reviews in the Neurosciences 32, 737–750. doi:10.1515/revneuro-2020-0146.

73. Zhao, Y., Zhang, Y., Meng, S., Chen, B., Dong, X., Guo, X., Guo, F., Zhang, R., Cui, H., and Li, S. (2023). Effects of S-Adenosylmethionine on Cognition in Animals and Humans: A Systematic Review and Meta-Analysis of Randomized Controlled Trials. J Alzheimers Dis 94, S267–s287. 10.3233/jad-221076.

74. Futschek, I.E., Schernhammer, E., Haslacher, H., Stögmann, E., and Lehrner, J. (2023). Homocysteine - A predictor for five year-mortality in patients with subjective cognitive decline, mild cognitive impairment and Alzheimer’s dementia. Exp Gerontol 172, 112045. 10.1016/j.exger.2022.112045.

75. Parkhitko, A.A., Wang, L., Filine, E., Jouandin, P., Leshchiner, D., Binari, R., Asara, J.M., Rabinowitz, J.D., and Perrimon, N. (2021). A genetic model of methionine restriction extends *Drosophila* health- and lifespan. Proceedings of the National Academy of Sciences 118, e2110387118. doi:10.1073/pnas.2110387118.

76. Hung, C.C., Chang, C.C., Huang, C.W., Nouchi, R., and Cheng, C.H. (2022). Gut microbiota in patients with Alzheimer’s disease spectrum: a systematic review and meta-analysis. Aging (Albany NY) 14, 477–496. 10.18632/aging.203826.

77. Saji, N., Niida, S., Murotani, K., Hisada, T., Tsuduki, T., Sugimoto, T., Kimura, A., Toba, K., and Sakurai, T. (2019). Analysis of the relationship between the gut microbiome and dementia: a cross-sectional study conducted in Japan. Scientific Reports 9, 1008. 10.1038/s41598-018-38218-7.

78. Mateo, D., Marquès, M., Domingo, J.L., and Torrente, M. (2024). Influence of gut microbiota on the development of most prevalent neurodegenerative dementias and the potential effect of probiotics in elderly: A scoping review. American Journal of Medical Genetics Part B: Neuropsychiatric Genetics 195, e32959. 10.1002/ajmg.b.32959.

79. Wu, L., Han, Y., Zheng, Z., Peng, G., Liu, P., Yue, S., Zhu, S., Chen, J., Lv, H., Shao, L., et al. (2021). Altered Gut Microbial Metabolites in Amnestic Mild Cognitive Impairment and Alzheimer’s Disease: Signals in Host–Microbe Interplay. Nutrients 13, 228.

80. Zheng, J., Zheng, S.-J., Cai, W.-J., Yu, L., Yuan, B.-F., and Feng, Y.-Q. (2019). Stable isotope labeling combined with liquid chromatography-tandem mass spectrometry for comprehensive analysis of short-chain fatty acids. Analytica Chimica Acta 1070, 51–59. 10.1016/j.aca.2019.04.021.

81. Oblak, A.L., Lin, P.B., Kotredes, K.P., Pandey, R.S., Garceau, D., Williams, H.M., Uyar, A., O’Rourke, R., O’Rourke, S., Ingraham, C., et al. (2021). Comprehensive Evaluation of the 5XFAD Mouse Model for Preclinical Testing Applications: A MODEL-AD Study. Front Aging Neurosci 13, 713726. 10.3389/fnagi.2021.713726.

82. Sterniczuk, R., Antle, M.C., Laferla, F.M., and Dyck, R.H. (2010). Characterization of the 3xTg-AD mouse model of Alzheimer’s disease: part 2. Behavioral and cognitive changes. Brain Res 1348, 149–155. 10.1016/j.brainres.2010.06.011.

83. Differding, M.K., Doyon, M., Bouchard, L., Perron, P., Guérin, R., Asselin, C., Massé, E., Hivert, M.F., and Mueller, N.T. (2020). Potential interaction between timing of infant complementary feeding and breastfeeding duration in determination of early childhood gut microbiota composition and BMI. Pediatr Obes 15, e12642. 10.1111/ijpo.12642.

84. Chénard, T., Malick, M., Dubé, J., and Massé, E. (2020). The influence of blood on the human gut microbiome. BMC Microbiol 20, 44. 10.1186/s12866-020-01724-8.

85. Yu, Z., García-González, R., Schanbacher, F.L., and Morrison, M. (2008). Evaluations of different hypervariable regions of archaeal 16S rRNA genes in profiling of methanogens by Archaea-specific PCR and denaturing gradient gel electrophoresis. Appl Environ Microbiol 74, 889–893. 10.1128/aem.00684-07.

86. Yu, Z., and Morrison, M. (2004). Improved extraction of PCR-quality community DNA from digesta and fecal samples. Biotechniques 36, 808–812. 10.2144/04365st04.

87. Yu, Z., Yu, M., and Morrison, M. (2006). Improved serial analysis of V1 ribosomal sequence tags (SARST-V1) provides a rapid, comprehensive, sequence-based characterization of bacterial diversity and community composition. Environ Microbiol 8, 603–611. 10.1111/j.1462-2920.2005.00933.x.

88. Caporaso, J.G., Lauber, C.L., Walters, W.A., Berg-Lyons, D., Huntley, J., Fierer, N., Owens, S.M., Betley, J., Fraser, L., Bauer, M., et al. (2012). Ultra-high-throughput microbial community analysis on the Illumina HiSeq and MiSeq platforms. Isme j 6, 1621–1624. 10.1038/ismej.2012.8.

89. Kozich, J.J., Westcott, S.L., Baxter, N.T., Highlander, S.K., and Schloss, P.D. (2013). Development of a dual-index sequencing strategy and curation pipeline for analyzing amplicon sequence data on the MiSeq Illumina sequencing platform. Appl Environ Microbiol 79, 5112–5120. 10.1128/aem.01043-13.

90. Callahan, B.J., McMurdie, P.J., and Holmes, S.P. (2017). Exact sequence variants should replace operational taxonomic units in marker-gene data analysis. Isme j 11, 2639–2643. 10.1038/ismej.2017.119.

91. Paulson JN, O.N., Braccia DJ, Wagner J, Talukder H, Pop M, Bravo HC (2013). metagenomeSeq: Statistical analysis for sparse high-throughput sequencing. http://www.cbcb.umd.edu/software/metagenomeSeq.

92. Paulson, J.N., Stine, O.C., Bravo, H.C., and Pop, M. (2013). Differential abundance analysis for microbial marker-gene surveys. Nature Methods 10, 1200–1202. 10.1038/nmeth.2658.

93. McMurdie, P.J., and Holmes, S. (2013). phyloseq: An R Package for Reproducible Interactive Analysis and Graphics of Microbiome Census Data. PLOS ONE 8, e61217. 10.1371/journal.pone.0061217.

94. Bolyen, E., Rideout, J.R., Dillon, M.R., Bokulich, N.A., Abnet, C.C., Al-Ghalith, G.A., Alexander, H., Alm, E.J., Arumugam, M., Asnicar, F., et al. (2019). Reproducible, interactive, scalable and extensible microbiome data science using QIIME 2. Nature Biotechnology 37, 852–857. 10.1038/s41587-019-0209-9.

95. Lin, H., and Peddada, S.D. (2020). Analysis of microbial compositions: a review of normalization and differential abundance analysis. NPJ Biofilms Microbiomes 6, 60. 10.1038/s41522-020-00160-w.

96. Law, C.W., Zeglinski, K., Dong, X., Alhamdoosh, M., Smyth, G.K., and Ritchie, M.E. (2020). A guide to creating design matrices for gene expression experiments. F1000Res *9*, 1444. 10.12688/f1000research.27893.1.

97. Ritchie, M.E., Phipson, B., Wu, D., Hu, Y., Law, C.W., Shi, W., and Smyth, G.K. (2015). limma powers differential expression analyses for RNA-sequencing and microarray studies. Nucleic Acids Res 43, e47. 10.1093/nar/gkv007.

98. Douglas, G.M., Maffei, V.J., Zaneveld, J.R., Yurgel, S.N., Brown, J.R., Taylor, C.M., Huttenhower, C., and Langille, M.G.I. (2020). PICRUSt2 for prediction of metagenome functions. Nat Biotechnol 38, 685–688. 10.1038/s41587-020-0548-6.

99. Yang, C., Mai, J., Cao, X., Burberry, A., Cominelli, F., and Zhang, L. (2023). ggpicrust2: an R package for PICRUSt2 predicted functional profile analysis and visualization. Bioinformatics 39. 10.1093/bioinformatics/btad470.

