## Supplementary figures and images for "Ketogenic interventions prevent alterations of the gut microbiome in transgenic Alzheimer’s Disease mice"

### Supplemental figure 1

#### Supplementary figure 1

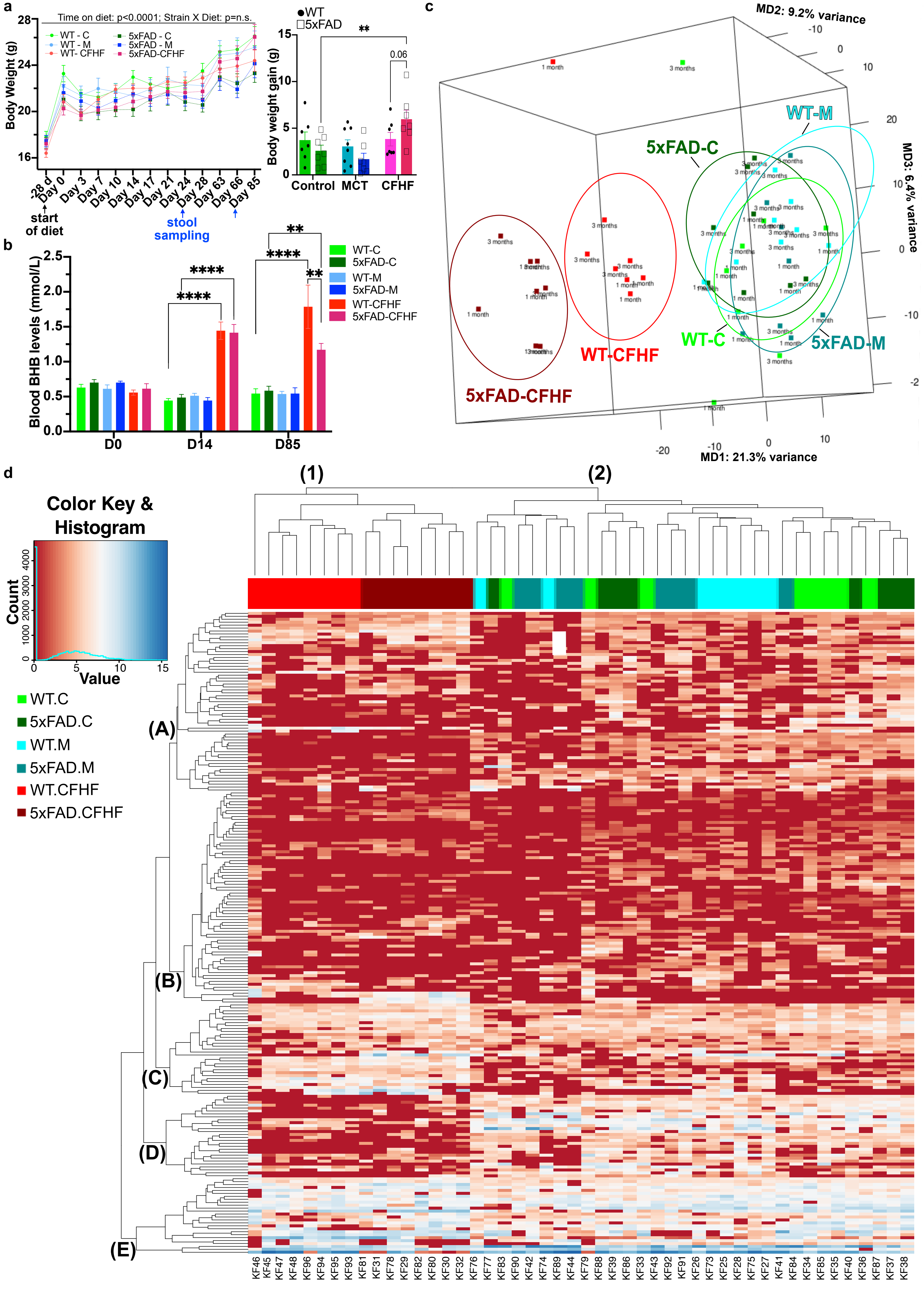

### Supplemental figure 2

#### Supplementary figure 2

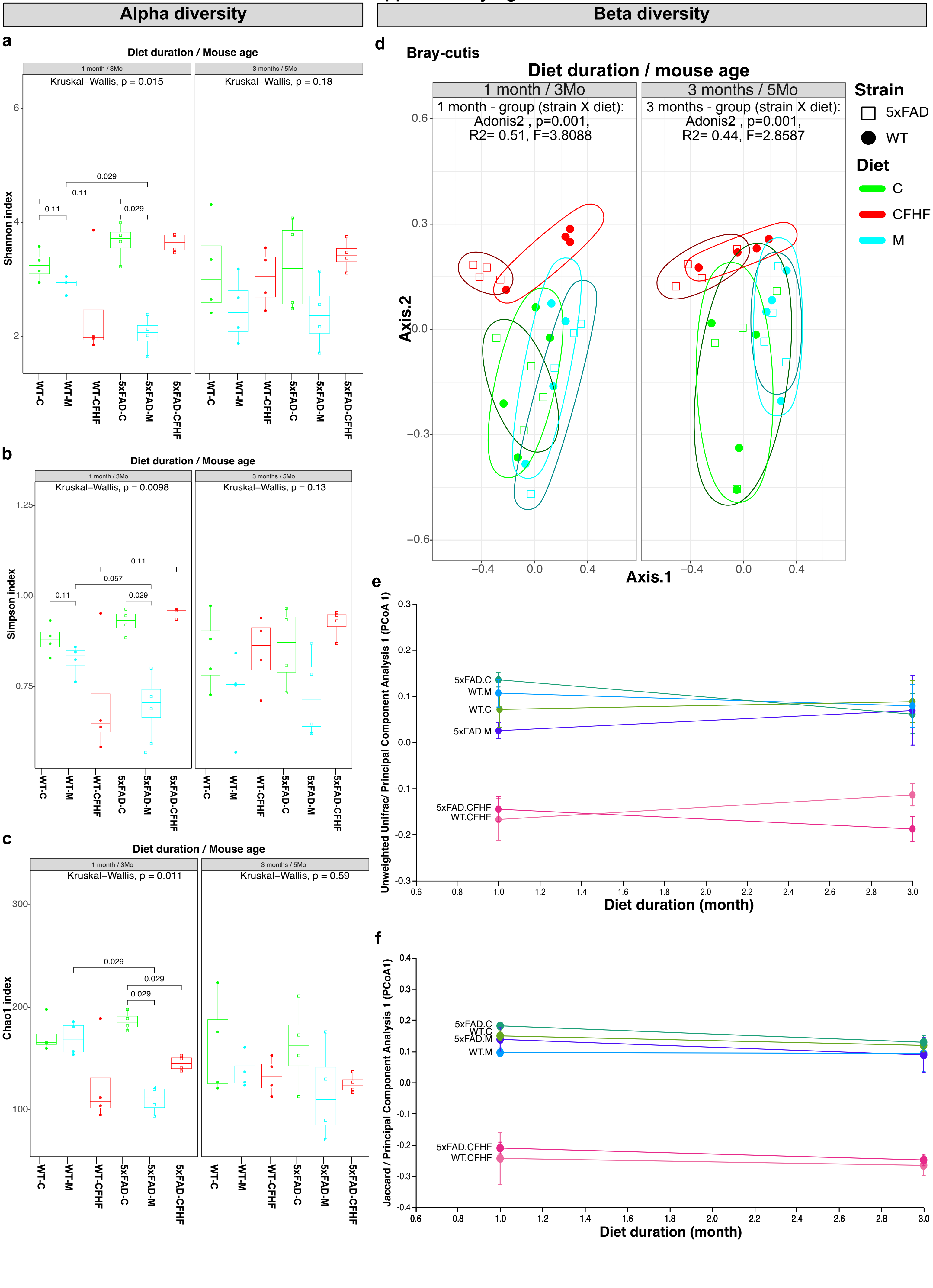

### Supplemental figure 3

**Suppplementary figure 3**

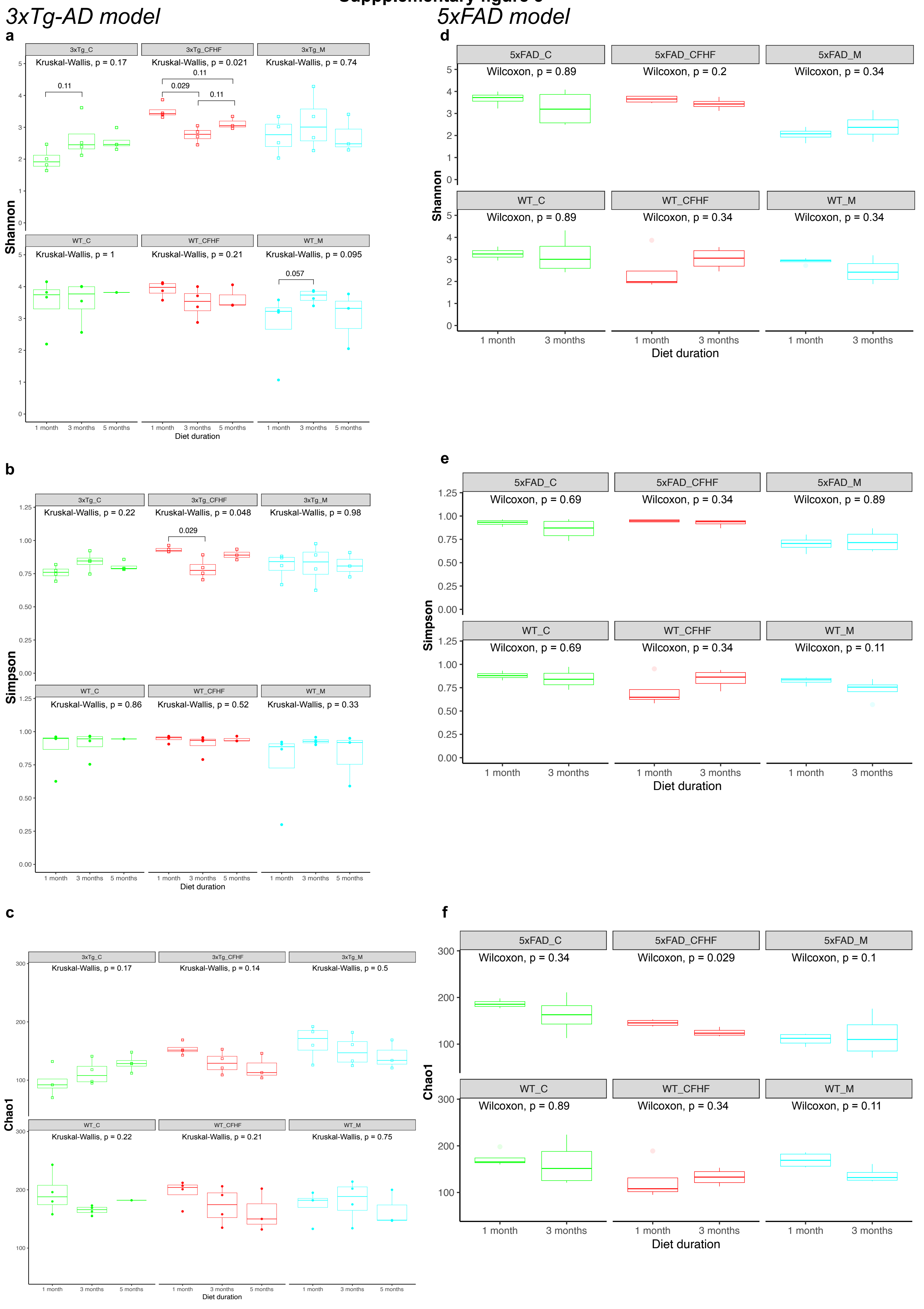

### Supplemental figure 4

#### Supplementary Figure 4

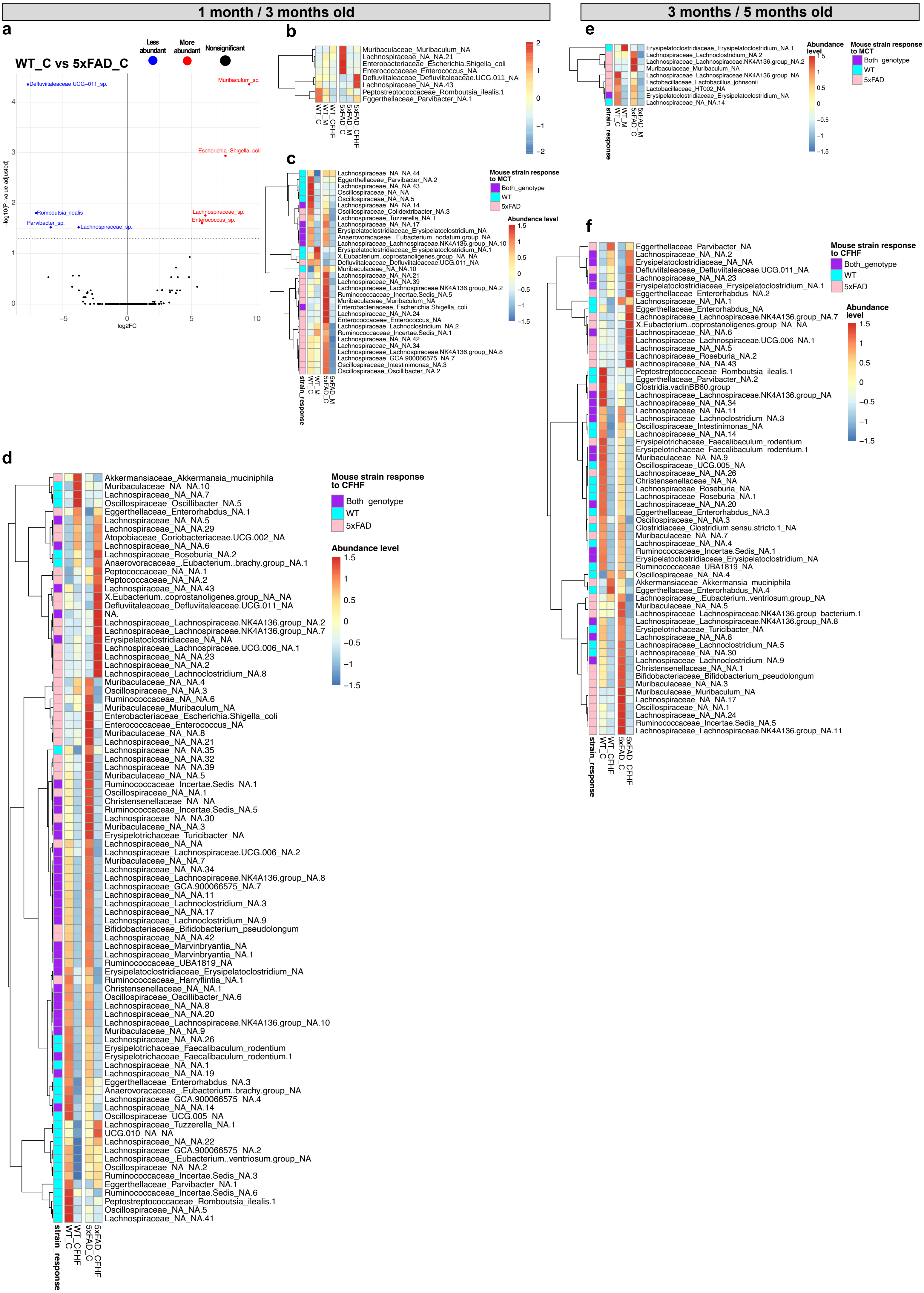

### Supplemental figure 5

#### Supplementary figure 5

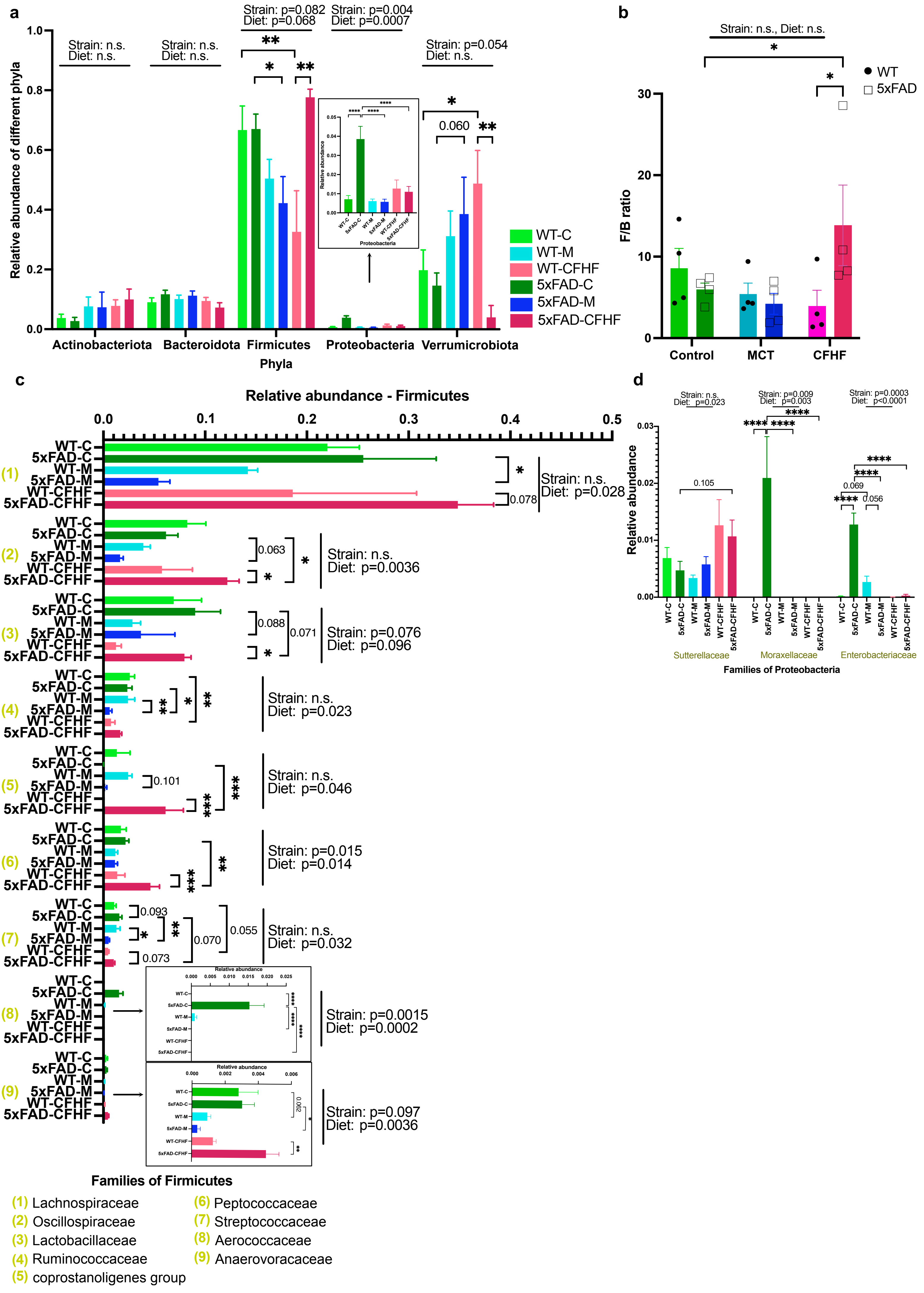

### Supplemental figure 6

#### Supplementary figure 6

a

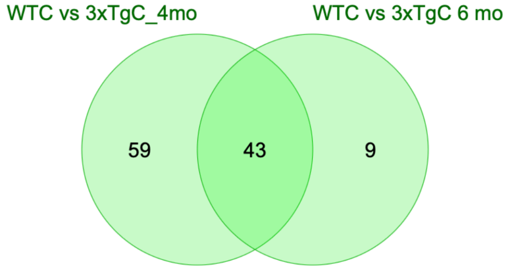


b

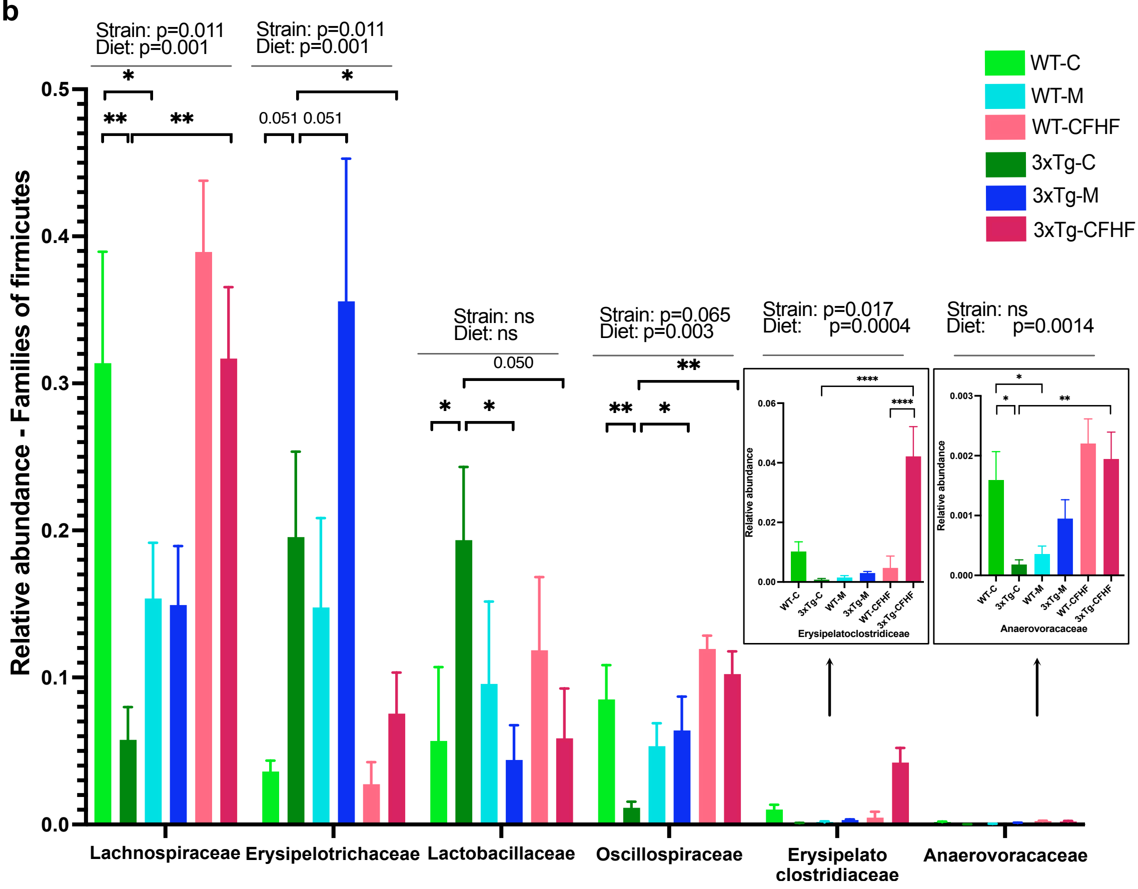


c

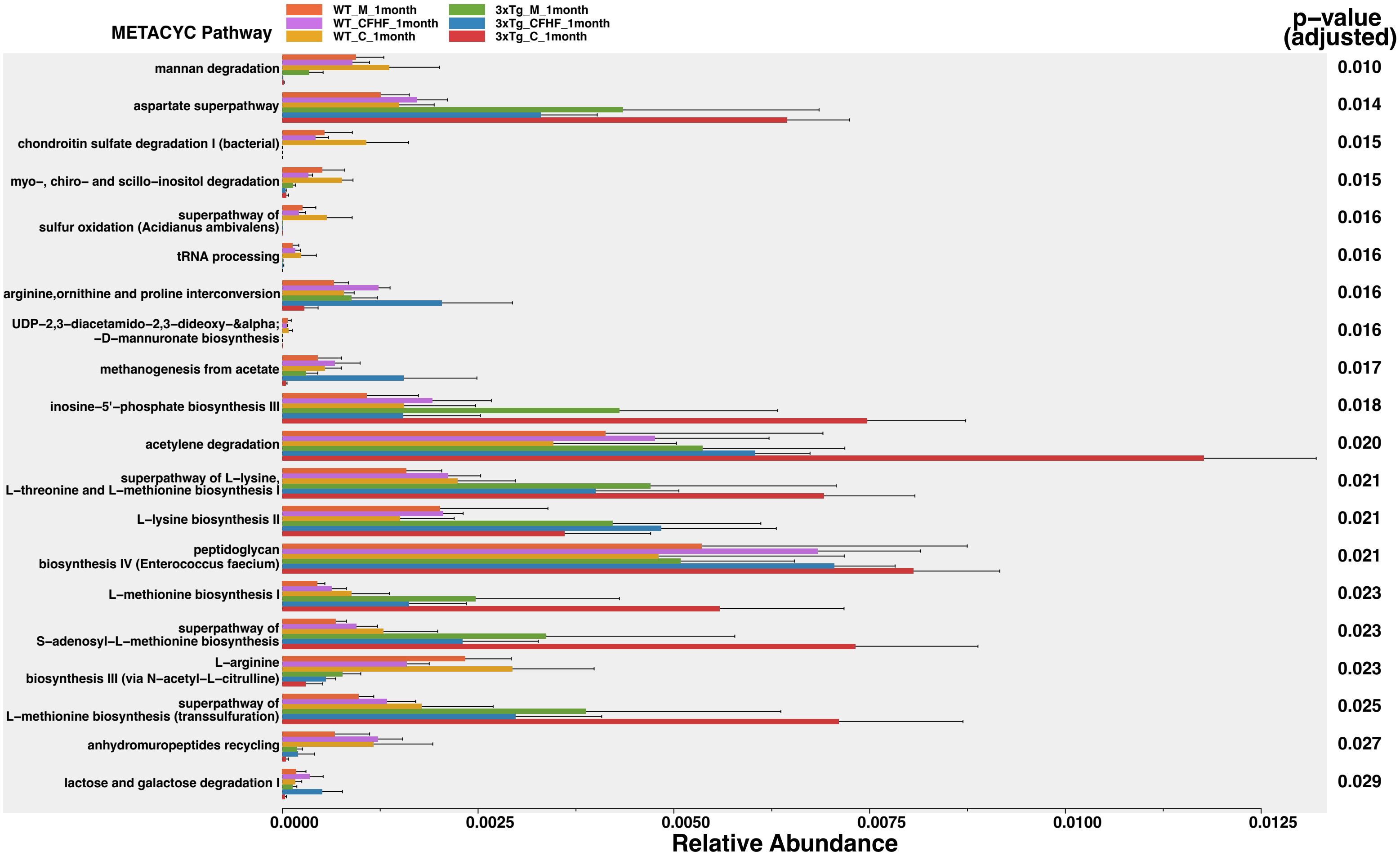

### Supplemental figure 7

##### Supplementary figure 7

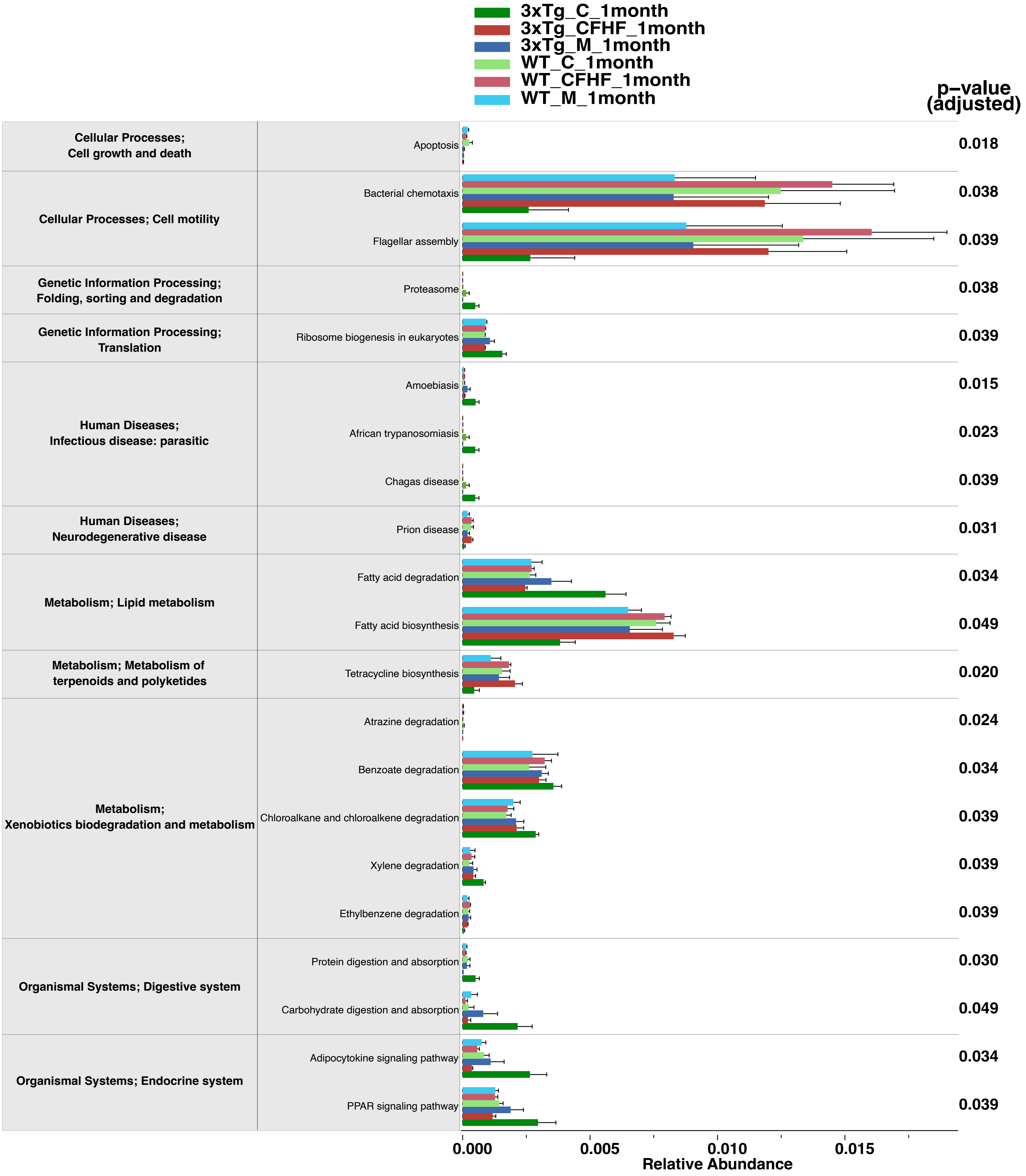

### Supplemental figure 8

**Supplementary figure 8**

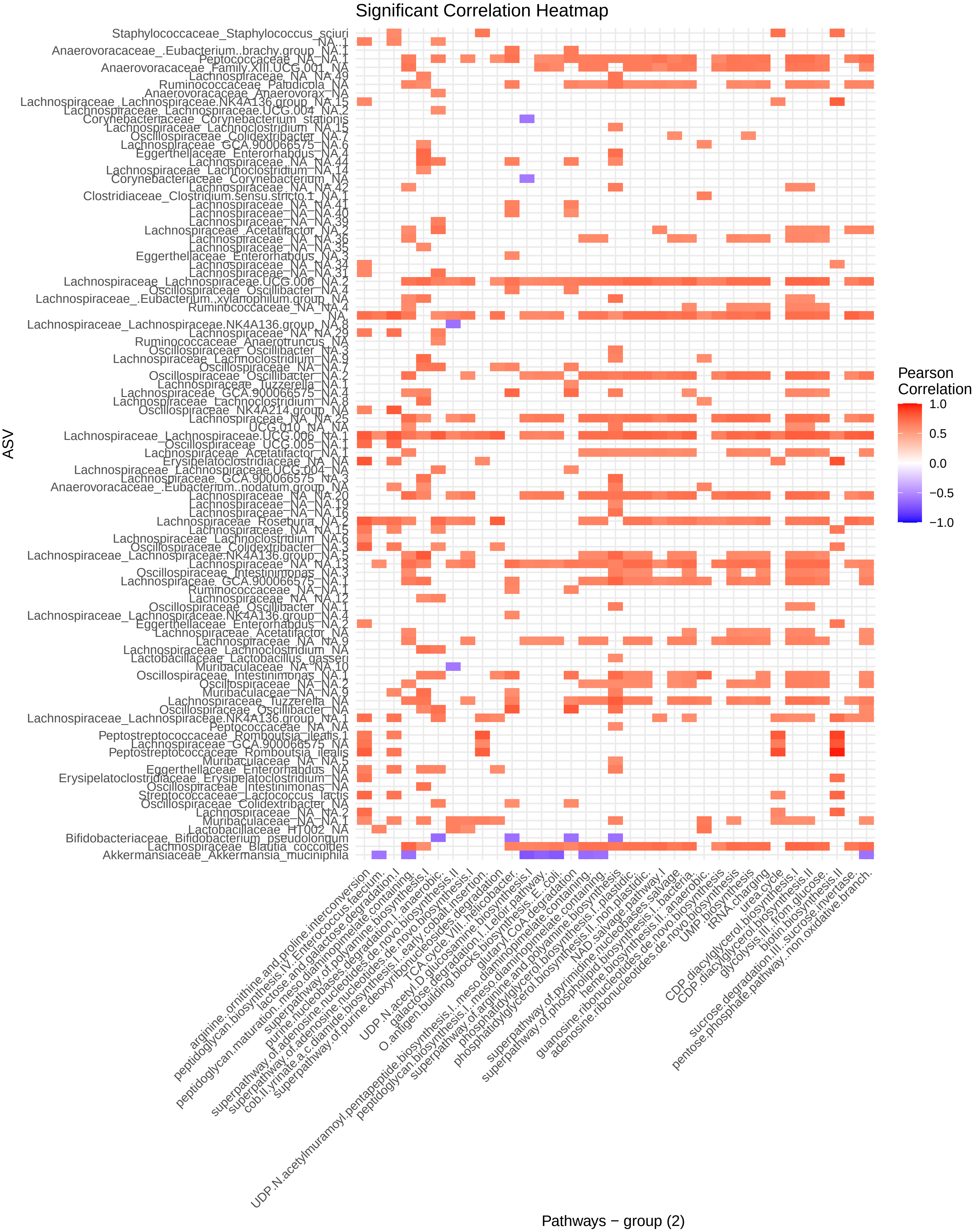
